# DNA six-way junction conformations and their use in a 2D square lattice

**DOI:** 10.1101/2025.04.09.648031

**Authors:** Alexander J. Baten, Hunter G. Mason, Remi Veneziano, William P. Bricker

**Affiliations:** Department of Chemical and Biological Engineering, University of New Mexico, Albuquerque, New Mexico; School of System Biology, George Mason University, Fairfax, Virginia; Department of Bioengineering, George Mason University, Fairfax, Virginia

## Abstract

In this paper, we investigate the conformational landscape of the DNA six-way junction (6WJ), a higher-order extension of the DNA four-way junction (4WJ), to determine preferred structural isomers. The 6WJ allows for unique junction-level topologies, and could be used to create novel DNA nanostructures, but the conformational landscape for the 6WJ is much more complex than the 4WJ. Our proposed conformational landscape for the 6WJ includes eight unique structural motifs, including an unstacked motif, five distinct stacked motifs, and two twisted motifs, and we estimate that there are forty structural isomers for the 6WJ, compared to only three for the 4WJ. To gain insight into these conformations, we perform all-atom molecular dynamics (MD) simulations on fourteen of the structural isomers. Our analysis shows that each 6WJ motif can be distinguished by its duplex stacking angles, and only a few of the 6WJ isomers have favorable free energies, which include the planar parallel isomers, the twisted isomers, and one of the orthogonal isomers. Lastly, to confirm that these stacked 6WJ motifs could be used to create larger-scale DNA nanostructures, we design a 2D square lattice using four orthogonal motifs connected in a 2×2 arrangement. Experimental characterization of this 2D lattice shows folding into the correct size and shape, further confirmed by AFM images of the DNA tiles, and an MD simulation that shows a slight twist of the lattice structure from orthogonality. This is the first 2D tile assembly built using an orthogonal junction-level topology for DNA nanotechnology, allowing a smaller mesh size than is possible with the 4WJ architecture.

## Introduction

DNA nanotechnology takes advantage of complementary Watson-Crick base-pairing to produce pre-programmed self-assembling nanostructures, where DNA nanostructures can be constructed from periodic networks of geometrical shapes as originally envisioned by Seeman.^1^ These nanostructures can be constructed from connected multi-arm junctions, the most recognizable of which are the three-way or four-way (3WJ/4WJ) ^1,2^ junctions inspired by intermediates of genetic recombination. ^3^ The development of DNA origami,^4^ a method involving the careful sequence design of a long ssDNA “scaffold” strand and many smaller ssDNA “staple” strands that self-assemble in one step, was a huge advancement in DNA nanotechnology. Assuming a well defined sequence, the folding of the scaffold and staple ssDNA strands would occur in one step with few errors, eliminating a reliance on stoichiometric precision and long purification steps.^5^ Furthermore, the DNA origami method allows the precise positioning of molecular attachments and the creation of more complex nanostructures.

For many applications of DNA nanotechnology the base building block is the immobile 4WJ,^6,7^ which connects two double-stranded DNA (dsDNA) duplexes in a parallel orientation. The immobility of the junction locks each strand in place and prevents branch isomerization, and is dependent on the sequence symmetry at the junction and adjacent strands. In isolation the 4WJ has been extensively studied by experiments^8,9^ and theory,^10,11^ which have determined that this junction exists in three conformations:^12^ an unstacked open-X conformation, and two stacked anti-parallel isomers which prefer a right-handed twist angle near 60*^◦^*,^13^ although this is sequence dependent. When used as a building block for DNA nanotechnology, the simplest combination is a sequential chain of 4WJs called a double crossover (DX) tile.^14^ This basic DX motif has been integrated into more complex 3D shapes such as the 6-helix bundle (6HB) ^15^ and 8 helix-bundle (8HB) ^16^ motifs, as well as in 3D polyhedral structures,^17,18^ and in 2D^19,20^ and 3D^21^ lattices. These structures can be used as building blocks to create giga-dalton or micron-scale DNA structures such as meta DNA (M-DNA), ^22^ self-limiting rings,^23^ the DNA pointer,^24^ and the Twisttower.^25^ All of the aforementioned DNA nanostructures have applications in areas such as photonics by creating nanowires ^26,27^ and synthetic energy transporting systems,^28–32^ for targeted drug delivery,^33–35^ as elements in DNA computing architectures,^36^ to promote self-assembly of other nanoparticles,^37^ and in bioimaging.^34,35,38^

Besides the 4WJ, alternative building blocks for DNA nanotechnology have been proposed and researched, such as a triple crossover (TX) junction made of two consecutive offset 4WJs,^39^ paranemic crossover (PX) tiles which contain crossovers at every possible point along the sequence and are constructed from a sequence of parallel as opposed to anti-parallel 4WJs,^40^ the PX’s topoisomer JX,^41^ anti/meso-junctions,^42^ the double multi-arm junction (DMAJ) consisting of multiple DX junctions side by side, ^43^ and the layered crossovers (LX)^44^ consisting of two DX crossovers connected and stacked on top of each other. These alternative building blocks have also been used to create larger structures such as a 2D crystal, ^45^ a 3D meso-junction lattice,^46^ conductive nano-wires,^27^ and 2D/3D tessellations.^47^ Other unique building blocks that have been explored include the tensegrity triangle ^48^ which has been utilized to make 3D crystal lattices,^49^ and a variety of N-arm branched junctions with 5, 6, 8, and 12 arms.^50–52^ Only the 6-arm and 8-arm branched junctions have been explored for incorporation into larger structures as a 2D lattice,^43^ with multi-armed junctions also being proposed to be used in 3D crystal lattices.^50,51^ Although many of these alternative building blocks have been explored using theory and experiment, none of them have been as successful and widely incorporated in DNA nanotechnology as the 4WJ and the DX motif.

The advent of DNA origami helped to catalyze the development of new tools to assist in the design and quantification of novel DNA junctions and structures. A variety of computational tools and simulations have facilitated these developments by allowing the direct visualization and study of the structural dynamics of DNA nanostructures at or near the atomic level, a resolution not accessible to experiment. Computational tools for routing the scaffold strand, such as caDNAno,^53^ DAEDALUS,^54^ Tiamat,^55^ ATHENA,^56^ and Adenita^57^ have assisted in the design and sequence routing of novel DNA origami structures. In addition, these DNA nanostructures can be simulated at the atomic level with all-atom molecular dynamics (MD), although time-steps must be at atomically-relevant time scales (∼1 fs), so simulations at biologically-relevant time scales (µsec to sec or longer) are difficult to impossible to simulate, especially with larger systems.^58^ One solution to the scaling problem is to use coarse-grained molecular dynamics,^59^ available through programs such as OxDNA, ^60^ which is helpful for simulating larger structures and for longer time scales. In this study we use all-atom MD for the individual junction simulations.

Much of DNA nanotechnology has centered around DX tiles and multi-dimensional extensions based on the 4WJ motif, with the ability to create 1D,^19^ 2D,^20^ and 3D^61^ structures. However, this building block contains only four duplex arms and therefore a relatively limited number of conformational isomers^12,62^ and sequence possibilities.^11,63^ Therefore, multi-arm branched junctions such as the six-way junction (6WJ) which is studied here present an opportunity for a diversified set of conformational isomers which can be used to create novel DNA nanostructures and lattices useful for a wider variety of applications. Previous studies of the 6WJ involved the quantification of duplex stacking of an open 6-armed star, with results confirming stacking on two adjacent arms.^50^ Another study creates our planar parallel motif experimentally by integrating it into a 1D and 2D scaffold.^43^ However, each of these studies only explored a portion of the potential 6WJ conformational landscape. Therefore, we have identified and explored the conformational landscape of the 6WJ motif in detail, by identifying all of the possible stacked 6WJ isomers generated from one sequence. Each structure has been simulated using all-atom molecular dynamics simulations, with a detailed analysis of its structural geometry and energetics. The isolated 6WJ has also been experimentally folded and characterized, confirming that it folds into a monodisperse population and that all strands are incorporated. To provide evidence that the 6WJ motif can be used in an extended lattice configuration, we choose an orthogonal 6WJ isomer and show that a 2×2 tile based on this motif can be routed and experimentally characterized. In future work, we will attempt to create multiple novel lattice configurations based on different structural isomers of the 6WJ.

## Results and Discussion

We explore multi-arm DNA junction space starting with the six-way junction (6WJ), which is the subsequent even-armed branched junction in complexity following the 4WJ. Even-armed junctions are useful as they can form duplex-stacking domains of integer values (i.e., a 4WJ can form two stacked duplexes and a 6WJ can form three stacked duplexes). The 6WJ has been studied previously with the original application being DNA crystal lattices. This study however was limited to attempting to identify the structural features of the predominately forming structure in solution. ^50^ The 6WJ has more recently been studied for tile-based self assembly,^43^ however this study only looked at a small portion of the potential conformational landscape of the 6WJ, not studying all of the building blocks that could be formed. In order to apply the 6WJ as a potential building block to larger DNA nanostructures, we have adapted a nomenclature analogous to that between the 4WJ and the DX motif, and as such the term triple crossover (TX) motif could be used for extensions of the 6WJ, as three DNA strands crossover at the same base-pair position. To not cause confusion with the aforementioned TX tiles^39^ which do not have three strands crossing over at the same position, we rationalize that a proposed conformational change from Open to the Planar Parallel motif displayed in Figure 1 is analogous to the conformational change of the unstacked 4WJ to the anti-parallel stacked DX motif, and assert that a continuous crossover at the same branch point between three duplexes should be considered a triple crossover. An example of the uniqueness of the 6WJ is shown with the hypothesized Top Orthogonal motif, which can be reached by further isomerization from the Planar Parallel motif, allowing for orthogonal directional changes in a DNA nanotechnology object at the junction level. The chosen sequence and experimental characterization of this isolated 6WJ is discussed in the following section.

**Figure 1:**
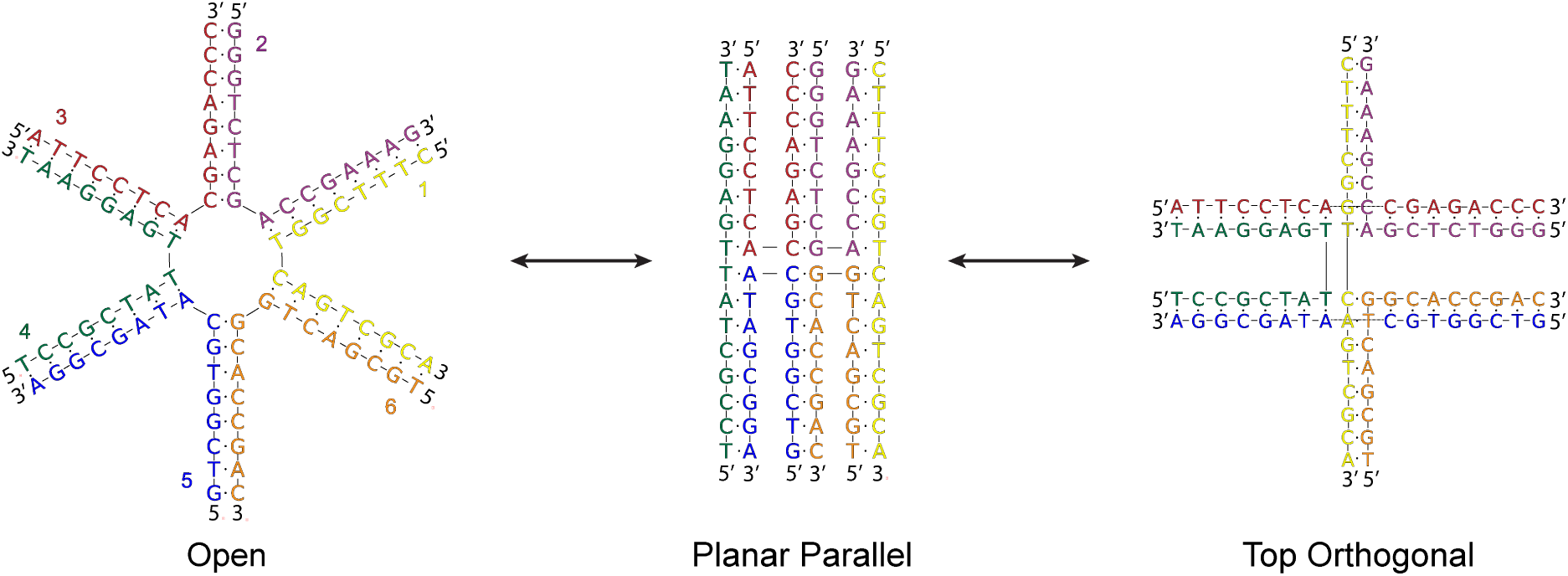
Proposed isomerization pathways in the six-way junction motif. A theoretical conformational pathway of the DNA six-way junction (6WJ) starting from the Open (left) to the Planar Parallel (center) and finally to the Top Orthogonal (right) motif. The Open motif of the 6WJ is analogous to the Open motif of the 4WJ with two additional strands. The Planar Parallel motif is analogous to the anti-parallel DX motif, corresponding to three duplexes stacking instead of two. The Top Orthogonal motif is formed from the Planar Parallel motif when one of the outside stacked duplexes migrates to the top of the other DNA strands, and the other four arms each rotate 90*^◦^* and form new duplex stacking partners. The DNA strands are labeled according to color on the Open motif as yellow (1), purple (2), red (3), green (4), blue (5), and orange (6), and this labelling is consistent throughout the manuscript for the six-way junction.

### Sequence and folding of a DNA six-way junction

To synthesize the DNA six-way junction motif, we first designed the sequences for the 6 oligonucleotides (16 nucleotides each) required for the assembly as described in Table S2 and as seen pictorially in Figure 1. The sequence design was done using the software Tiamat 2^55^ to create orthogonal sequences and avoid the risk of mispairing that would cause biproduct formation. The sequences designed were then validated with BLAST using the blastn algorithm (with the word size set at 7).^64^ Once validated, the sequences were acquired and mixed at an equimolar concentration in the folding buffer prior to be annealed overnight and run on an agarose gel to validate the proper assembly. The results presented in Figure 2A show that the folding of the DNA six-way junction leads to the formation of a monodisperse population of nanoparticles as indicated by the single band visible on the gel. To further validate the correct formation of the junction, we included multiple FRET pairs in the motif design. One donor dye (fluorescein) was placed on the 5’-ends of strand numbers 1 and 5 (Table S2) and one acceptor dye was placed on the 5’-ends of strand numbers 2, 3, and 6. This organization of FRET dyes allow us to cover the entire surface of the nanoparticles and the distances at which they have been placed and estimated on our model are compatible with FRET. The results presented in Figure 2B-C show that energy transfer is occurring in all FRET pair conformations tested, validating the formation of the nanoparticles and the incorporation of all strands. Moreover, the results observed are compatible with some of the potential conformations as predicted with our models. However, these results are not sufficient to inform on the preferred conformation(s), as the potential conformational landscape as described in the next section is too complex. Nevertheless, the experimental characterization of the DNA six-way junction here confirms that we have a folded structure, and we use molecular modeling in the next few sections to try to determine which folded 6WJ conformations are most likely. The experimental setup for the sequence and folding of the DNA six-way junction is discussed in detail in the *Experimental Methods* section.

**Figure 2:**
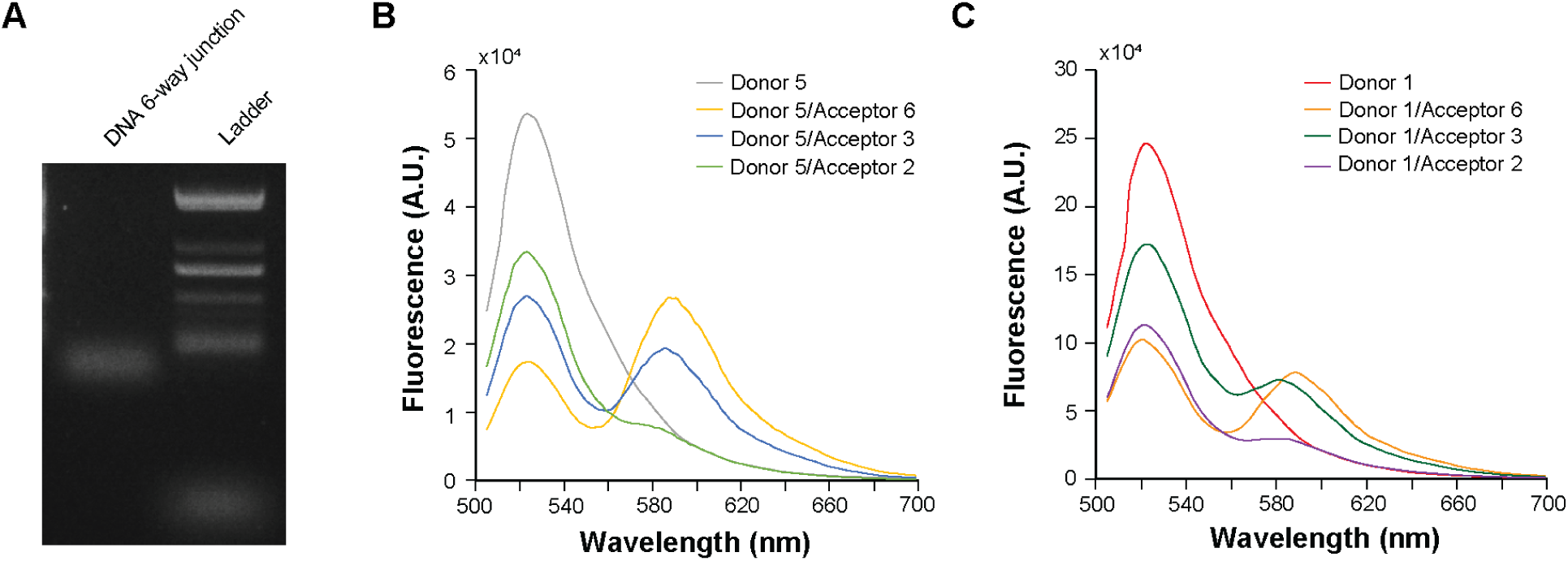
Sequence and experimental characterization of a DNA six-way junction. (A) Agarose gel electrophoresis validating the formation of the DNA six-way junction. (B) Fluorescence resonance energy transfer using strand 5 as the donor and 3 different strands (2, 3, and 6) as the acceptor. (C) Fluorescence resonance energy transfer using strand 1 as the donor and 3 different strands (2, 3, and 6) as acceptor.

### Conformational landscape of DNA six-way junction isomers

The potential conformational landscape of a DNA six-way junction is much more complicated than a DNA four-way junction, which has two unique motifs (stacked and unstacked), and five unique isomers for each junction sequence including open, DX,^14^ and PX^40^ junctions. For the DNA six-way junction, eight unique structural motifs are identified, including one unstacked motif and seven stacked motifs (Figure 3). The unstacked motif is labeled as **Open**, and is analogous to the open motif seen in the DNA four-way junction except with two additional arms.^65^ As seen in numerous studies involving the four-way junction, the open form is not energetically preferred over the stacked forms due to electrostatic and dispersive effects,^65,66^ which was also alluded to in previous studies of the 6WJ, ^43,50^ so we anticipate that the open form of the six-way junction will not be energetically favored over conformations where duplexes are allowed to stack. The first stacked motif occurs when the **Open** six-way junction folds into a planar arrangement of three stacked duplexes, referred to as planar parallel and abbreviated as **PP**. The planar parallel motif can also be thought of two side-by-side stacked four-way junctions, and the difference in minor/major groove widths of the central duplex creates a slightly convex shape when the minor groove of the central duplex at the triple crossover is viewed from above.

**Figure 3:**
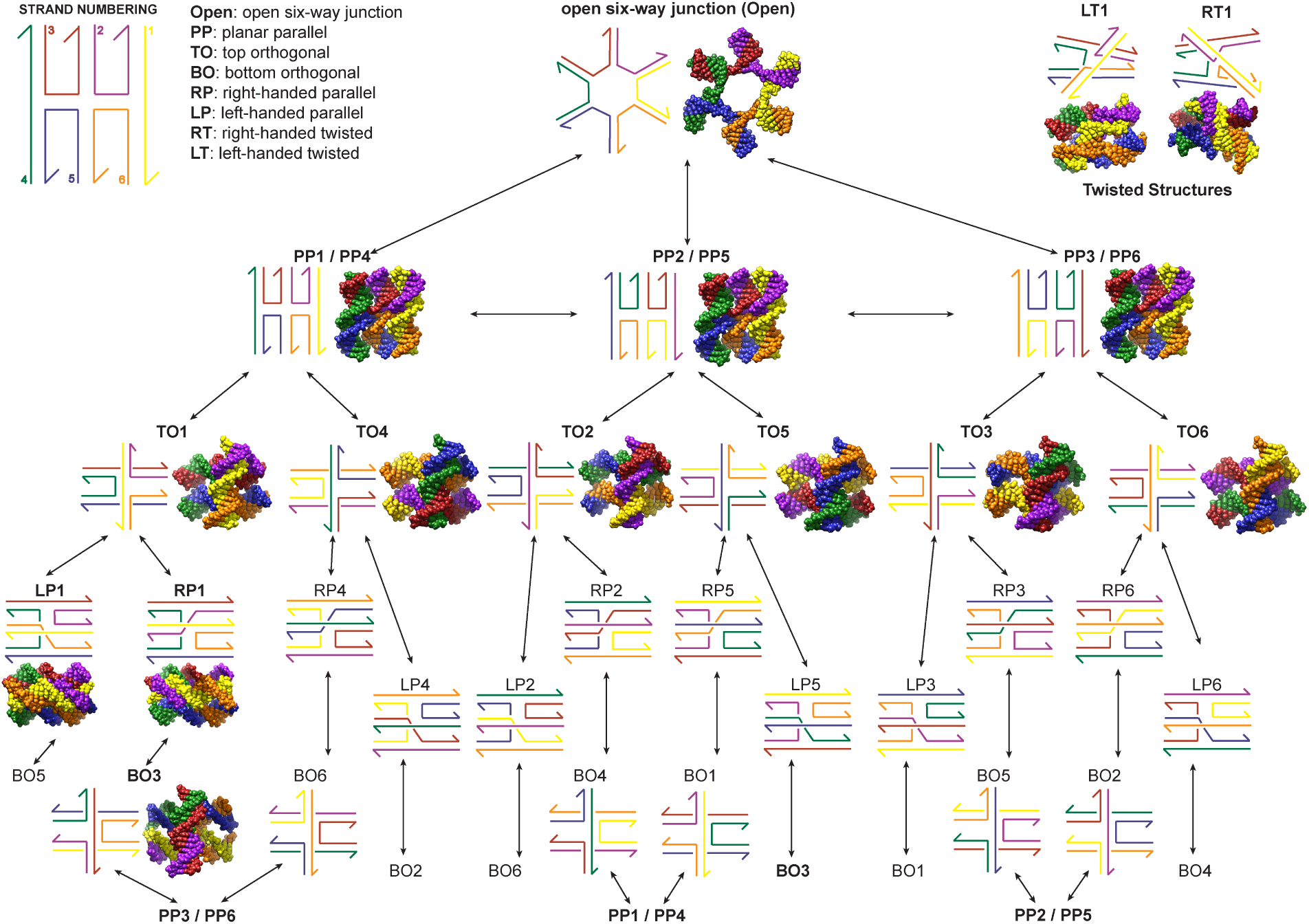
Potential conformational motifs and isomers of a DNA six-way junction. The DNA strands are labeled and colored as strand 1 (yellow), strand 2 (purple), strand 3 (red), strand 4 (green), strand 5 (blue), and strand 6 (orange). Eight unique structural motifs for the DNA six-way junction are identified, including one unstacked structure (**Open**), five stacked structures (**PP**: planar parallel, **TO**: top orthogonal, **BO**: bottom orthogonal, **RP**: right-handed parallel, **LP**: left-handed parallel), and two twisted structures (**RT**: right-handed twisted, **LT**: left-handed twisted). Arrows between conformations indicate likely pathways between structural motifs. The number following the structural motif identifier indicates the unique isomeric structure based on sequence (e.g. TO1 is the top orthogonal structure with strand 1 in the orthogonal position). A molecular model of each unique structural motif is shown corresponding to the all-atom molecular dynamics simulations performed in this study.

The remainder of the stacked motifs can be split into orthogonal stacked structures, parallel stacked structures, and the stacked and twisted structures. The orthogonal stacked structures are reached directly from the planar parallel motif and are referred to as top orthogonal (**TO**) and bottom orthogonal (**BO**), where the direction indicates whether the orthogonal duplex has folded on the top or bottom of the original planar parallel structure. To achieve these orthogonal structures, the two left-most duplexes in the planar parallel structure must switch stacking partners (i.e., similar to four-way junction isomer switching), and the right-most duplex will end up on the top or bottom of the re-folded structure. The parallel stacked structures can be reached from the orthogonal stacked structures when the orthogonal duplex rotates into a parallel arrangement with the other two duplexes. These parallel stacked structures are referred to as right-handed parallel (**RP**) and left-handed parallel (**LP**), where the handedness refers to the direction of rotation of the orthogonal duplex to achieve each motif directly from the top orthogonal motif. Lastly, the stacked and twisted motifs, referred to as right-handed twisted (**RT**) and left-handed twisted (**LT**), are intermediate between the orthogonal stacked and parallel stacked motifs, where the handedness refers to the direction of duplex twist around the long-axis of the structure, as well as whether they are folding towards the **RP** or **LP** structures. These twisted motifs are likely to be sampled during any conformational shift between an orthogonal stacked and parallel stacked motif.

While there is only one sequential isomer of the **Open** motif, each of the stacked motifs can fold into multiple isomeric structures. The top orthogonal and bottom orthogonal motifs each have six isomeric forms which are labeled according to which sequence strand is the orthogonal strand. In addition, the right-handed parallel and left-handed parallel motifs each have six isomeric forms which are labeled according to which orthogonal strand has rotated from the top orthogonal motif. The right-handed twisted and left-handed twisted motifs additionally have six isomeric forms each as these structures are intermediate between the orthogonal and parallel stacked motifs, although only two out of twelve are shown. The only exception here is the planar parallel motif which has only three unique isomeric forms due to rotational symmetry between **PP1/PP4** structures, the **PP2/PP5** structures, and the **PP3/PP6** structures. All in all, this provides a much more complicated conformational landscape for the DNA six-way junction as compared to the DNA four-way junction that includes eight unique structural motifs and a grand total of *forty* potential structural isomers, most of which are shown in Figure 3. (Only two out of the twelve possible twisted isomeric structures are shown.) In order to analyze these DNA six-way junction structures in greater detail, we built atomic models for each of the unique structural motifs, and ran and extensively analyzed all-atom molecular dynamics (MD) simulations for information regarding structure, dynamics, and energetics, all of which are elaborated in the following sections.

### Geometry of DNA 6WJs from all-atom molecular dynamics simulations

To begin our analysis of the isolated 6WJ, we built atomic models of twelve unique structural isomers as shown in Figure 3, including the **Open**, **PP1 / PP2 / PP3**, **TO1 / TO2 / TO3 / TO4 / TO5 / TO6**, **LP1**, **RP1**, **BO3**, **LT1**, and **RT1** motifs. These atomic models were solvated with explicit water and ions, and we performed three replicates of allatom molecular dynamics (MD) for 150 ns for each structural isomer, giving a total of 450 ns of simulation time for each model. Extensive details regarding how these atomic models were built and the all-atom MD simulation parameters is provided in the *Computational Methods* section.

### Duplex-duplex angles for individual 6WJ isomers from all-atom MD simulations

As we are interested in the stacked isomeric configurations of the 6WJ, a key structural identifier is the angle between duplex pairs during the all-atom MD simulations. The duplexes are defined from the 5’ to 3’ end of each strand, and are labeled as **d**_1_, **d**_2_, and **d**_3_, which are analogous to duplex strands 1 (yellow), 3 (red), and 5 (blue) in each motif’s isomer 1 conformation, as shown in Figure 3, except for the **BO** which corresponds to **BO3** in Figure 3. For the **TO** and **PP** isomers the duplexes contain the same numbering even though the strand colors will have switched. Snapshots from MD are extracted for each model, and the angles during the simulation between each of the duplex pairs was found using the dot product of the two normalized vectors corresponding to each duplex and it’s direction, as described further in the *Computational Methods* section. These angles between duplex pairs are shown in Figure 4 for each structural model, where it is apparent that each geometric motif has a distinct and recognizable set of duplex angles, although these angles are roughly the same between isomers of the same motif, such as between **TO** or **PP** isomers.

**Figure 4:**
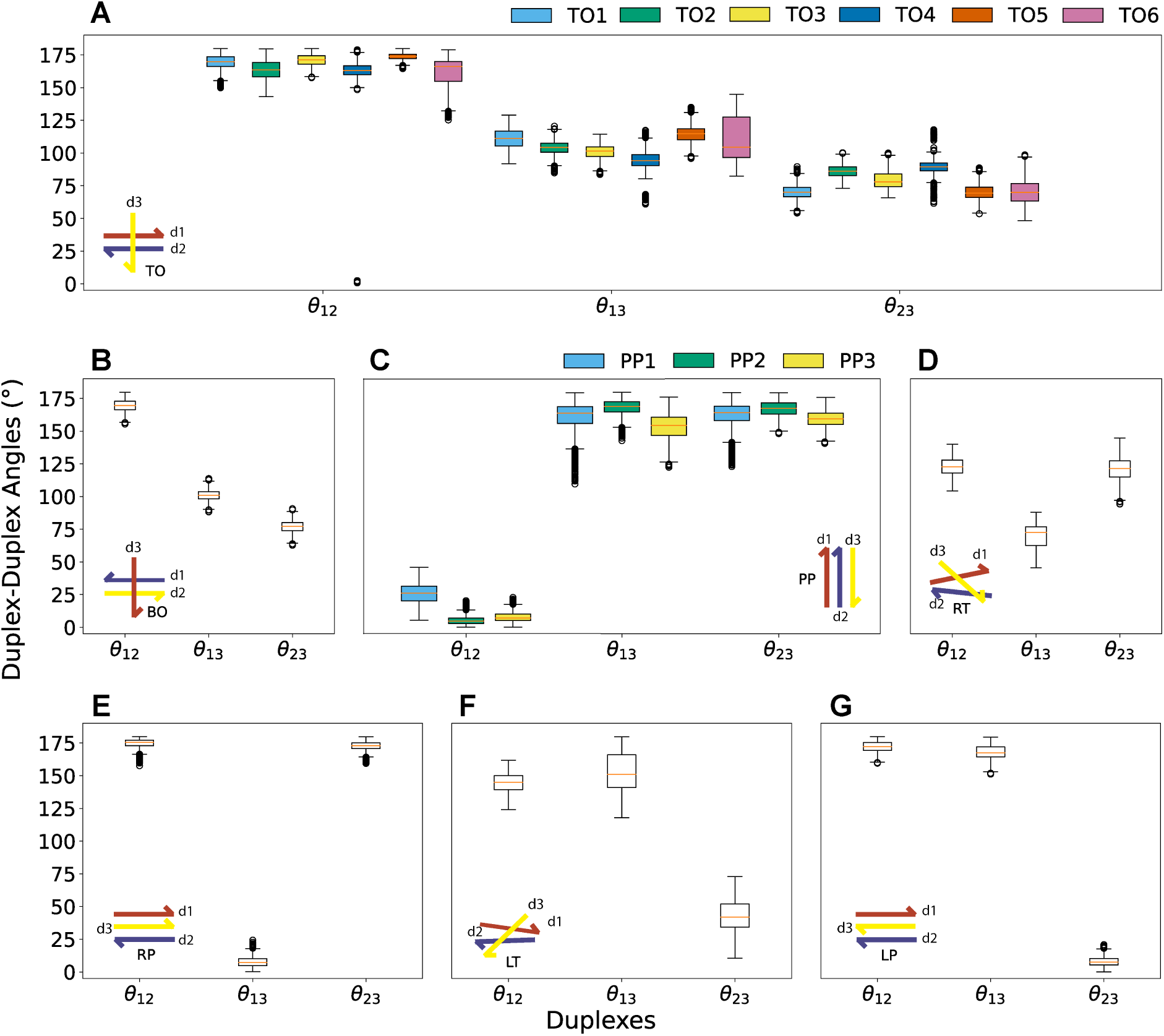
**Duplex stacking angles for all simulated 6WJ isomers.**Shown are boxplots of the duplex-duplex stacking angles (*^◦^*) between the first (**d**_1_), second (**d**_2_), and third (**d**_3_) duplexes for each motif-isomer combination simulated with all-atom molecular dynamics, where θ_12_ describes the angle between the **d**_1_ and **d**_2_, θ_13_ is the angle between the **d**_1_ and **d**_3_, and θ_23_ is the angle between the **d**_2_ and **d**_3_. The duplex-duplex stacking angles for (A) top orthogonal isomers **TO1**-**TO6**, (B) bottom orthogonal isomer **BO3**, (C) planar parallel isomers **PP1**, **PP2**, and **PP3**, (D) right-handed twisted isomer **RT1**, (E) right-handed parallel isomer **RP1**, (F) left-handed twisted isomer **LT1**, and (G) left-handed parallel isomer **LP1**. The whiskers represent the minimum and maximum values excluding outliers, while the top and bottom edges of the box contain data points in the 25th-75th percentile with the orange line indicating the median, and outliers are greater than 1.5 times the inter-quartile range.

The orthogonal motifs (**TO** and **BO**) tend to have one duplex pair with anti-parallel angles and the other two duplex pairs with nearly orthogonal angles. In Figure 4A, for **TO1** the angle between **d**_1_ and **d**_2_ (θ_12_) is nearly anti-parallel at 171*^◦^* and **d**_3_ is close to orthogonal for both **d**_1_ and **d**_2_ where θ_13_ = 111*^◦^* and θ_23_ = 70.3*^◦^*. The rest of the top orthogonal isomers (**TO2-6**) follow a similar pattern with θ_12_ nearly anti-parallel and θ_13_ and θ_23_ nearly orthogonal, with a slight difference in orthogonality between the isomers. Comparatively, in Figure 4B, the **BO3** motif exhibits very similar patterns to the **TO** motifs with θ_12_ nearly anti-parallel and θ_13_ and θ_23_ orthogonal, although the overall spread from the MD snapshots is tighter than for the **TO** isomers.

The parallel motifs (**PP**, **RP**, and **LP**) each have all three duplex pairs with nearly parallel or anti-parallel angles. In Figure 4C, the **PP** motifs tended to include parallel or anti-parallel duplex-duplex angles. For isomer **PP1**, the angle between **d**_1_ and **d**_2_ averaged 26.0*^◦^*, and this angle for **PP2** and **PP3** was much closer to parallel, at 5.33*^◦^*and 7.82*^◦^*, respectively. The other duplex-duplex angles were close to anti-parallel with θ_13_ = 160.4*^◦^* and θ_23_ = 162.3*^◦^* for **PP1**, and similar values for **PP2** and **PP3**. Similar to the **PP** motifs, all of the duplex-duplex angles for **RP1** were either parallel or anti-parallel where θ_12_ = 174.7*^◦^*, θ_13_ = 7.53*^◦^*, and θ_23_ = 172.8*^◦^*, as shown in Figure 4E. The left-handed version, **LP1** has similar parallel or anti-parallel average angles of θ_12_ = 172.2*^◦^*, θ_13_ = 168.1*^◦^*, and θ_23_ = 7.81*^◦^*, as shown in Figure 4G. These last two parallel motifs (**RP1** and **LP1**) have the tightest spread in duplex-duplex angles of all of the structures studied, indicating that they may be constrained in a local structural minima.

Lastly, the twisted motifs (**RT** and **LT**) have duplex-duplex angles that are generally skewed (neither parallel, anti-parallel, or orthogonal). As shown in Figure 4D, **RT1** has duplex-duplex angles of θ_12_ = 122.7*^◦^*, θ_13_ = 70.2*^◦^*, and θ_23_ = 120.9*^◦^*. Similarly, in Figure 4F, **LT1** has duplex-duplex angles of θ_12_ = 144.7*^◦^*, θ_13_ = 42.0*^◦^*, and θ_23_ = 152.6*^◦^*. These twisted motifs should be viewed as having intermediate angles between the orthogonal and parallel motifs excluding **PP**.

### Geometry and stability of DNA 6WJ motifs

The deviation of the duplex-duplex angles for each isomer indicates relatively stability (even if constrained to a local minima) and each has a distinct and identifiable set of duplex-duplex angles, additionally providing some insight into the flexibility or rigidity of each isomer.

The set of orthogonal isomers (**TO** and **BO**) all maintain similar duplex-duplex angles throughout the MD simulation that indicates that they are located at or near a local structural minima. The only exception is **TO6**, which displayed a large range of duplex-duplex values, potentially due to sequence-related effects at the junction. Despite this all orthogonal isomers maintain RMSD values of less than 10Å (Figures S1-S7), implying no large conformational changes during the simulation time. Contrary to their name, these isomers have angles that are slightly skewed from orthogonal which is due to the asymmetric atomic structure of dsDNA (i.e., the phosphate backbone of each DNA strand is not equidistant leading to the asymmetry between the minor and major groove side of the DNA duplex). This asymmetry causes a mismatch of the minor and major grooves of **d**_3_ (the orthogonal strand) and the anti-parallel strands (**d**_1_ and **d**_2_) when in contact, as the junction must minimize phosphate-phosphate backbone interactions between the strands. Due to these forced backbone interactions, the set of orthogonal motifs in isolation are not likely to be as energetically favorable as other isomers that can more easily minimize these phosphate-phosphate interactions. These orthogonal isomers can be readily identifiable from other motifs by their duplex-duplex angles, indicated by a anti-parallel θ_12_ angle, and nearly orthogonal θ_13_ and θ_23_ angles, where θ_13_ tends to skew obtusely and θ_23_ tends to skew acutely. Unfortunately, **TO** and **BO** cannot be identified from each other based solely on duplex-duplex angles.

The **PP** isomers exhibit flexible and relatively stable unique duplex-duplex angles, with the exception of **PP1** whose **d**_3_ angles contain a large number of outliers deviating from the mean, indicating that **d**_3_ may have moved slightly out-of-plane. Out-of-plane duplex-duplex angles are not unexpected in the isolated **PP** motif, since the isolated 4WJ prefers out-of-plane twist angles near 60*^◦^*,^13^ and the **PP** motif is essentially two side-by-side 4WJ motifs. Due to limitations on the MD simulation time-scales, it is unclear if this was the beginning of a conformational transition or just random fluctuations. The stability of these isomers is confirmed with the RMSD plots displayed in Figures S8-S10 that peak around 10Å, smaller than expected from a full conformational transition. All duplex-duplex angles between isomers maintained relatively similar means and spreads indicating that all **PP** motifs are identifiable by their duplex-duplex angles when compared to other motifs, specifically a parallel θ_12_ angle, and anti-parallel θ_13_ and θ_23_ angles. The remaining two parallel motifs (**RP** and **LP**) are essentially a 3-helix bundle displaying near planar duplex-duplex angles. The lack of deviation in each duplex-duplex box-plot alludes to both **RP1** and **LP1** existing in relatively unfavorable and constrained local minima, with little room for movement. While these duplex-duplex angles are either parallel or anti-parallel like the **PP**, all three parallel motifs can be distinctly identified. The **RP** motif has a parallel θ_13_ angle and anti-parallel θ_12_ and θ_23_ angles, and the **LP** motif has a parallel θ_23_ angle and anti-parallel θ_12_ and θ_13_ angles. Therefore, the three parallel motifs can be readily identified by which of the three duplex-duplex angles is parallel. RMSD plots for the parallel motif simulations are found in Figures S11 and S12.

As previously stated the two twisted motifs (**RT** and **LT**) do not have any duplex-duplex angles that are parallel, anti-parallel, or orthogonal, instead displaying skewed angles. This places the twisted motifs as intermediate conformations between the orthogonal motifs and the same-handed parallel motifs. Both twisted motifs also display the largest range of values for their angles indicating structural flexibility, as well as maintaining angles comparable to the favorable ∼60*^◦^* ^13^ twist conformation of the isolated Holliday junction. It should be noted however that **RT1** is more centered around the 60*^◦^* value, with the angles in **LT1** closer to parallel and anti-parallel, showing a distinct difference in structure based on handedness. Despite having the largest range for duplex-duplex angles, their RMSD (Figures S13 and S14) shows only marginal deviations, mostly for replicate two of **LT1**. The inherent structural flexibility implies that the twisted motifs are not potentially constrained to a local minima as seen for some of the orthogonal and parallel motifs, and are likely corresponding to a structural intermediates between these other motifs.

Each of the seven stacked 6WJ motifs present a unique set of duplex-duplex angles that can be differentiated with the exception of the two orthogonal configurations, and isomeric conformations of the same motif show small deviations from the average set of motif angles. Each isomer studied by the MD simulations is in a relatively stable geometry, supported by the consistent duplex-duplex angles and the small deviations from the RMSD plots in Figures S1-S14. We do note that each replicate of these isomers was only simulated for 150 ns, which is not enough simulation time to undergo a major conformational change. By quantifying the average structural components of each motif and isomer, we can identify how to integrate each motif into potential large-scale DNA nanostructures that utilize each motif as a building block, and we can start to identify which of the motifs and isomers may be more or less structurally and energetically favorable, as analyzed and discussed in the next section.

### Free energy calculations of isolated DNA 6WJ isomerization

The identification of which isolated 6WJ motifs are energetically preferred will allow the assessment of which large-scale DNA nanotechnology structures the 6WJ can be readily incorporated as a building block. We analyzed the free energies of isomerization using the Molecular Mechanics Poisson-Boltzmann Surface Area^67,68^ (MM-PBSA) method, which calculates the solvated free energy for each 6WJ isomer from all-atom MD by incorporating an implicit solvation potential derived from the Poisson-Boltzmann equation, discussed in detail in the *Computational Methods* section.

The MM-PBSA procedure was used to analyze the free energies of isomerization of the top orthogonal motifs **TO1**, **TO2**, **TO3**, **TO4**, **TO5**, and **TO6**, the bottom orthogonal motif **BO3**, the planar parallel motifs **PP1**, **PP2**, and **PP3**, the right-handed parallel motif **RP1**, the left-handed parallel motif **LP1**, the right-handed twisted motif **RT1**, and the left-handed twisted motif **LT1**. These calculations provide a quantitative understanding of the differences in free energy across this set of isomers, along with identifying the most energetically favorable isomer configuration. The Gibbs free energy (G) of the 6WJ isomers as calculated using MM-PBSA are shown in Figure 5, with the raw data provided in the SI in Table S1, broken down into the enthalpy (H), entropy (T S), and the Gibbs free energy (G), and box-plots for both H and T S are also provided in Figures S15 and S16.

**Figure 5:**
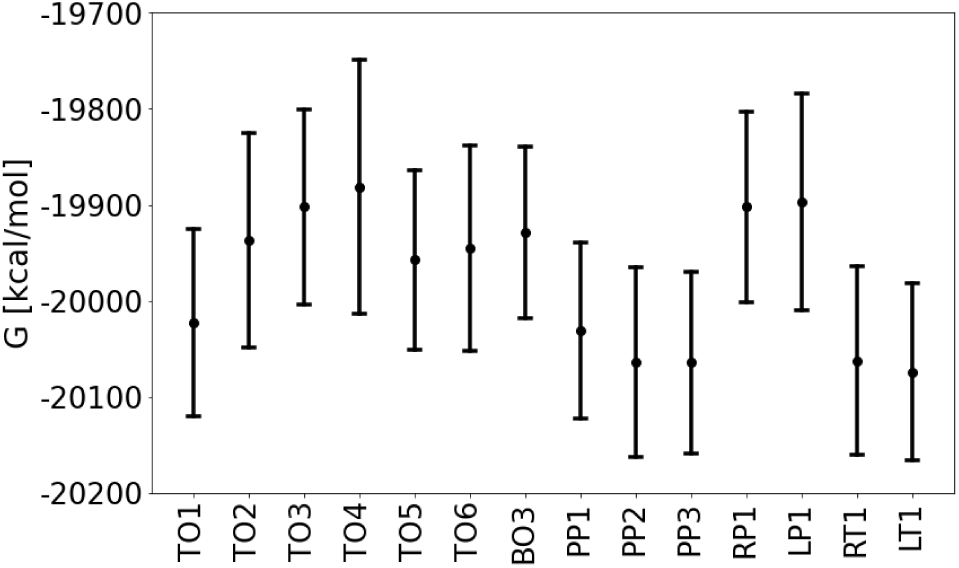
Free energies of isolated 6WJs. The calculated G of each simulated isomer using MM-PBSA including enthalpic and entropic contributions. The enthalpy values of each isomer were calculated using snapshots from all-atom molecular dynamics simulations taken every 100 ps of the last 125 ns for three replicates, for a total of 3753 samples for each isomer. The entropy of each isomer were calculated using snapshots taken every 500 ps from the last 125 ns for three replicate for a total of 750 samples for each isomer. The center indicates the mean value while the whiskers of the plot show the spread up to two standard deviations.

Unlike the previous trends identified between duplex-duplex angles of orthogonal motifs, there is a much larger set of free energy differences between the **TO** isomers. The lowest energy motif, **TO1**, had a value of −20023 kcal/mol, while the other top orthogonal isomers had more unfavorable G values of −19937 kcal/mol, −19902 kcal/mol, −19881 kcal/mol, −19957 kcal/mol, and −19945 kcal/mol for isomers 2 through 6 respectively. The bottom orthogonal **BO3** had a similarly unfavorable free energy value of −19929 kcal/mol. The parallel 6WJ motifs had a significant variation among isomers, with the planar parallel isomers **PP1**, **PP2**, and **PP3** exhibiting significantly lower free energies of −20031 kcal/mol, −20064 kcal/mol, and −20064 kcal/mol, respectively. The right- and left-handed parallel motifs simulated, **RP1** and **LP1**, were very unfavorable compared to the planar parallel motifs with −19902 kcal/mol and −19897 kcal/mol, respectively. Conversely, the right- and left-handed twisted motifs, **RT1** and **LT1**, had some of the most favorable free energies of −20062 kcal/mol and −20074 kcal/mol, respectively.

### Analysis of 6WJ free energies of isomerization and potential conformational pathways

The spread of the free energy values of each 6WJ isomer calculated, as shown in Figure 5, have varying degrees of overlap, indicating the extent of free energy values from snapshots in the all-atom MD simulations. We compare this free energy overlap as well as the ΔG (relative free energies of isomerization) between adjacent isomeric conformations, to determine the most favorable 6WJ motifs and isomers and potential routes for 6WJ conformational changes as hypothesized in Figure 3.

First, we discuss potential conformational changes between the planar parallel and orthogonal motifs, as the open motif is likely to initially stack as a planar parallel motif. As an example, **TO1** and **TO4** both experience some energetic overlap with **PP1**, which are directly connected via a relatively simple conformational change. The mean free energy of isomerization for **PP1** to transform into either top orthogonal isomer is ΔG = 8.2 kcal/mol and ΔG = 150 kcal/mol for **TO1** and **TO4** with both favoring **PP1**. Despite both being unfavorable transitions, a conformational change is possible, although a transformation to **TO1** is much more likely than to **TO4** as the free energy of isomerization is significantly less. Comparatively, the transition from **PP2** to either **TO2** or **TO5** has a free energy of isomerization of ΔG = 127 kcal/mol and ΔG = 107 kcal/mol, respectively, displaying a largely unfavorable transition. Finally, a conformational change between **PP3** and it’s adjacent conformational forms, **TO3** and **TO6**, is similarly considered unlikely due to a large free energy of isomerization of ΔG = 119 kcal/mol and ΔG = 196 kcal/mol, respectively. Similarly to most of the **TO** motifs the conformational change between **PP3** and **BO3** is also unfavorable with a free energy of isomerization of ΔG = 136 kcal/mol. The major difference between the isomers of the same motif is the sequence ordering in the core of the stacked 6WJ which implies that this core sequence plays a significant role in the preferred folding and potential conformational pathways.

From the free energy differences we hypothesize that the twisted conformations, **RT** and **LT**, can be reached from the orthogonal conformations, **TO** or **BO**, or directly from the planar parallel conformation. With this in mind, a transition from **TO1** to either **LT1** or **RT1** could occur depending on the handedness of the skew angle of the top strand. Both of the twisted conformations, **RT1** and **LT1**, have lower relative free energies than **TO1**, with a free energy of isomerization of ΔG = −38.3 kcal/mol for **RT1** and ΔG = −51.4 kcal/mol for **LT1**, making this a very favorable transition into either twisted conformation from the top orthogonal motif. As an alternate pathway, the twisted conformations have a free energy of isomerization of −31.12 kcal/mol and −43.24 kcal/mol for **RT1** and **LT1** transitioning from **PP1**, providing an additional pathway to form these motifs which may bypass forming an orthogonal structure completely. Conformational transitions from the twisted motifs, **RT1** and **LT1** to the parallel motifs, **RP1** and **LP1**, is considered unlikely due to an unfavorable free energies of isomerization of ΔG = 160 kcal/mol from **RT1** to **RP1** and ΔG = 177.2 kcal/mol from **LT1** to **LP1**. This leads us to conclude that the formation of isolated **RP1** or **LP1** conformations in solution to be unlikely or transient.

Overall, two of the planar parallel motifs (**PP2** and **PP3**) and both twisted motifs (**RT1** and **LT1**) have the lowest calculated free energy values, making these isomers the most likely to fold for the isolated 6WJ. This is somewhat unsurprising as both of these motifs have similarities to the preferred geometry of the isolated 4WJ, as the planar parallel motif is essentially two side-by-side 4WJs, and the twisted motif shares the preferred twist angle of near 60*^◦^* as seen in the isolated 4WJ. The other planar parallel motif **PP1** and the orthogonal motif **TO1** may additionally fold under specific conditions, but the other isomers simulated are not likely to fold due to unfavorable relative free energies. It should be noted that simulations of all possible isomers as presented in Figure 3 have not been completed due to computational constraints, and isomers of the same motif have already been shown to have vastly different free energies. In addition, these simulations were performed for one sequence of an isolated 6WJ, and it has been shown previously that the core sequence of an isolated 4WJ strongly influences its folding behavior. ^10,11^ Even with a core sequence change for the isolated 6WJ, we expect that the planar parallel and twisted conformations will be most favorable, although this is untested and presents a near impossible combinatorial problem for simulation. How much of the free energy difference is the result of the proximity of Mg^2+^ ions or sequence dependency^1,69–73^ is also unknown. Lastly, MM-PBSA is a rather coarse-grained method for estimating free energies as it is not clear whether the ionic contribution to junction stability is fully comprehensive. Furthermore, MM-PBSA is a end-state method capturing no details of the energetic landscape between conformations. Therefore in future studies we could investigate preferred conformational pathways with replica exchange methods,^74^ thermodynamic integration,^75^ or adaptive biasing force methods^76^ incorporated into our procedure. Even with these caveats, it has been shown here that the isolated 6WJ will have preferred stacking configurations, and the MM-PBSA calculations provide results that are fairly consistent with our expectations.

### Base-pair step parameters reflect isomer properties

Base-pair step (BPS) deviations can be interpreted as markers for structural changes needed to optimize energetic favorability in each isomer. Therefore, we expect the largest deviations to occur at the junction as each base-pair adjusts into a more favorable conformation. This can be seen for the **TO1**, **PP1**, and **LT1** isomers in Figure 6A-C where the BPS deviations for each duplex are displayed similar to the orientation of the atomic model of each motif. The three terminal BPS are removed from the plot as we are uninterested in quantifying end-fraying effects. The bases on each “duplex” correlate to the BPS of the continuous strand, with the fifth BPS from the 5’ end of each duplex representing the junction BPS. In general, the BPS far from the junction contain few deviations, but notably we can see that **TO1** and **PP1** tend to have more BPS deviations away from the junction as they are deforming to spatially accommodate the other stacked duplexes. For example, the negative deviation in twist for the **TO1** AC BPS in duplex 1 (red) corresponds to untwisting of the duplex where the minor groove needs to expand to minimize unfavorable interactions between the minor groove and the backbone of duplex 3 (yellow). Most of the other orthogonal motifs contain similar deviations of twist around the junction for similar reasons as the **TO1** isomer as seen in Figures S17-S22, where the top orthogonal duplex needs to deform to avoid the phosphate backbone atoms on the bottom duplexes.

**Figure 6:**
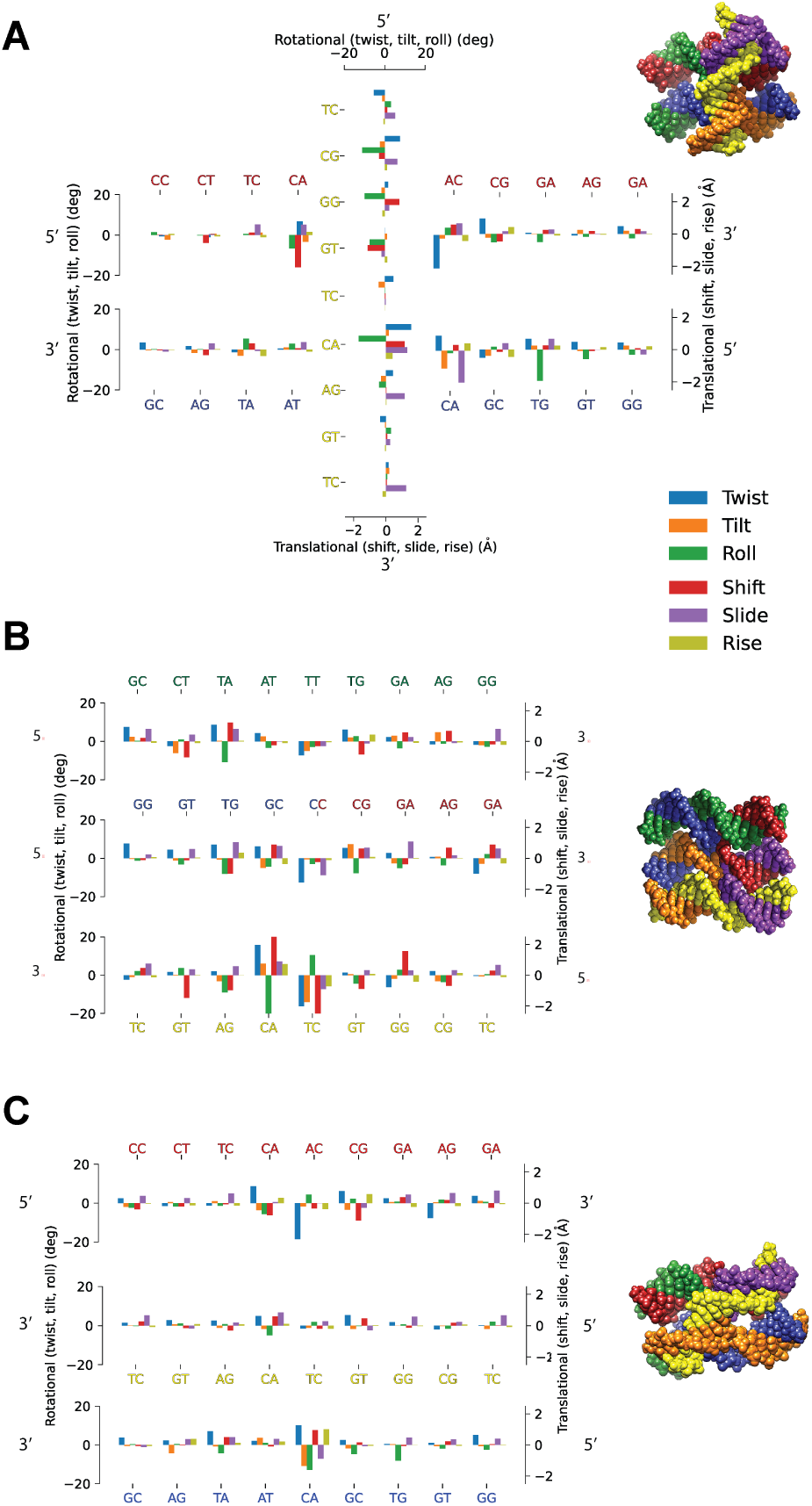
A comparison of conformational dynamics of the TO1, PP1, and LT1 isomers. The base-pair step (BPS) deviations of the (A) **TO1**, (B) **PP1**, and (C) **LT1** isomers compared to a reference B-DNA duplex are shown accompanied by a corresponding atomic model. BPS deviations in the twist, tilt, roll, shift, slide, and rise parameters are all indicated by color and both positive and negative deviations for each parameter are shown. For each BPS deviation plot the 5’ and 3’ on each side show the direction and BPS on the continuous strand.

Even though these BPS deviations can be indicators of favorable conformational adjustments at the junction, large amounts of deviations at and away from the junction can imply an unfavorable geometry due to excessive deformation. Out of all of the **TO** motifs, **TO1** has the most favorable energetics, as well as the smallest deviations away from the junction on the first and second duplex. The least favorable **TO** motifs are **TO3** and **TO4**, which have some of the largest BPS deviations at and away from the junction, indicating major structural deformation. These major deviations of BPS away from the junction are indicative of conformations not typically seen in the average B-DNA duplex, and should correlate to a less favorable duplex structure and isomeric conformation.

Comparatively, the more energetically favorable isomers such as the **PP1** motif and **LT1** have most of the larger BPS deviations centered at the junction and fewer away from the junction as seen in Figure 6B-C. The numerous BPS deviations throughout each of the duplexes in **PP1** are from adjustments made to minimize the forced backbone-backbone interactions of the parallel duplexes. The 4WJ can easily minimize these interactions by rotating the duplexes to a 60*^◦^* angle, but in the case of the **PP** motif it is not obvious how the three duplexes should rotate to accommodate each other. The other two **PP** isomers show a similar trend to **PP1** with BPS deviations throughout each duplex (Figures S25 and S26). Of all of the isomers, **LT1** has the smallest overall BPS deviations throughout all three of its duplexes (Figure 6C), which is unsurprising since it is also the most energetically favorable isomer tested. The twisted conformation allows each duplex to maintain a near 60*^◦^*angle with each other, which is known to be the most favorable configuration in the stacked 4WJ. The **RT1** isomer shares similar structural deformation characteristics with **LT1** and both twisted isomers have BPS deviation data in Figures S27 and S28.

Among the least favorable motifs are the two parallel motifs, **LP1** and **RP1**, which have large BPS deviations near the junction but smaller BPS deviations away from the junction (Figures S29 and S30). Of all the motifs, the parallel motifs have the most crowded junction space, clearly requiring major structural deformation to maintain their shape. This forced stacking of all three duplexes in the parallel motifs are a contributing factor to their unfavorable energetics.

### Sequence and folding of a DNA square lattice structure based on the top orthogonal 6WJ motif

Following the successful folding of the DNA six-way junction motif, we further investigated whether it could be used to assemble larger tiles, and focus our efforts here on a repeated TO motif. A repeated TO motif can be used to create a 2D square lattice, since the top strands are oriented orthogonally to the bottom strands. The isolated TO motif also showed fairly favorable free energies in our simulations, indicating that it might be amenable to incorporation into a larger scaffolded structure. We combined four of the TO junctions to create a 2 by 2 tile with an estimated size of 12 by 12 nm, and designed the sequences using Tiamat 2 as described earlier for the 6WJ.^55^ The sequence orthogonality was again validated using BLAST.^64^ For this 2 by 2 tile, we designed two variants, one with a long central single strand (80 nucleotides long) and one with the central strand split into two shorter strands as indicated in Table S4. These two variants use either 13 or 14 total strands for the single central strand variant and the two central strand variant, respectively. Similar to the isolated 6WJ we mixed the different oligonucleotides at equimolar concentrations in our folding buffer and annealed them overnight. Interestingly, the electrophoresis results presented in Figure 7A show a monodisperse population for the variant with a single central strand, while the variant with the split central strand shows bi-product that could correspond to a dimer (or multimer) formation (see Figure S31).

**Figure 7:**
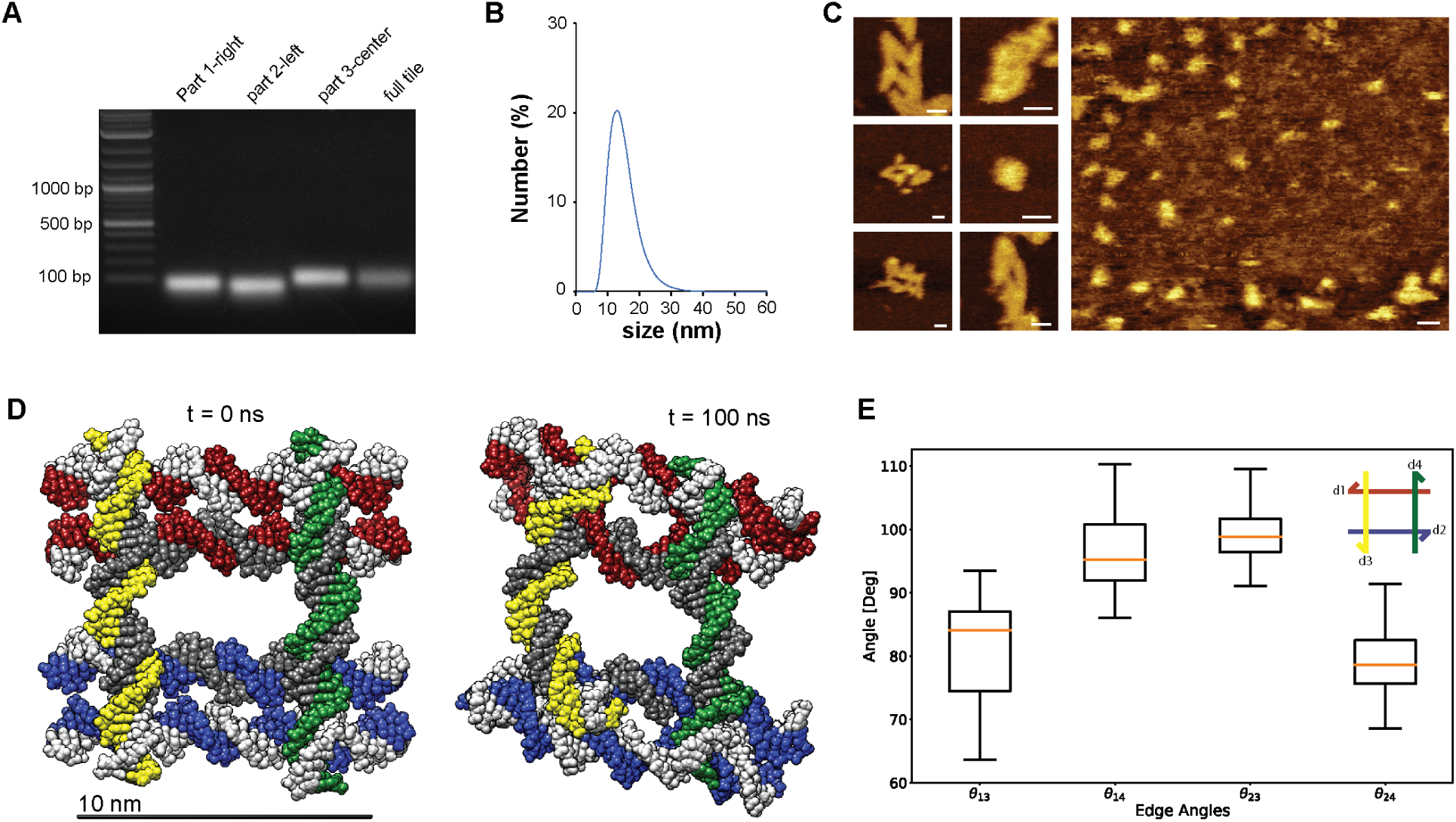
Folding and characterization of a 2D tile assembly based on four repeated TO motifs. (A) Agarose gel electrophoresis shows the sequential assembly of the 2D tile including different numbers of strands for the central staple strand. (B) Characterization of the size of the 2D tile using dynamic light scattering. (C) Atomic force imaging of the 2D tile on a bare surfece (left insets) or using a poly-D-lysine coating (right panel). (D) A molecular model of the 2D tile assembly with one central staple strand shows an ideal starting conformation (left) and a more twisted conformation after 100 ns (right). (E) The average interior angles from the MD simulation are indicative of the twisted structures seen in the AFM images.

Based on these results, we continued working with the single central strand variant only. Our step-by-step folding of this variant confirmed the proper formation of a single population of nanoparticles with a band for the fully formed tile between 100 and 200 bp (for a 216 bp structure) as expected for a densely packed nanostructure. We further performed dynamic light scattering measurements and measured an hydrodynamic diameter of 13.5 nm, which is slightly larger but consistent with the theoretical tile size estimated to be 12 nm in length. The AFM images of the tile also confirm the overall size and shape of the motifs but also indicate their propensity to form aggregates, which might be due to the presence of multiple blunt ends on each tile. In addition, it is important to note the slight deformation of the motifs into structures that are slightly skewed from orthogonal. Following this, we setup and performed a 150 ns MD simulation on the 2D tile lattice as shown in Figure 7D, which we initially place in an ideal orthogonal conformation, but the skewed conformations as seen in our AFM results are also indicated in the MD simulation (right panel). The average interior angles of the 2D tile lattice are close to orthogonal but prefer a skewed angle as seen in Figure 7E. We find these results consistent with the isolated 6WJ simulations, where the top orthogonal motif preferred angles slightly skewed from orthogonal, and an overall preference to form into the twisted motif, where all angles are closer to 60*^◦^*. Overall, our results here confirm the proper assembly of the square lattice motif using the 6WJ folded in a repeated TO motif.

## Conclusions

The DNA six-way junction (6WJ), which is a higher-order extension of the four-way junction (4WJ), provides the opportunity for unique DNA topologies at the junction-level that can be applied to create novel DNA nanostructures. Previous studies on the 6WJ^43,50^ explored isolated configurations of the 6WJ, which we expand upon by proposing a full conformational landscape for stacked 6WJs. From experiment, we confirm that the isolated 6WJ folds into a monodisperse population using gel electrophoresis. A subsequent FRET experiment with multiple FRET pairs confirms the incorporation of all strands into the structure, but cannot inform on which stacked 6WJ conformation is preferred.

To determine the preferred conformations of the isolated 6WJ, we first develop a potential conformational map including all stacked 6WJ motifs and isomers. This conformational map includes an unstacked **Open** motif, five stacked motifs (**PP**: planar parallel, **TO**: top orthogonal, **BO**: bottom orthogonal, **RP**: right-handed parallel, **LP**: left-handed parallel), and two twisted motifs (**RT**: right-handed twisted, **LT**: left-handed twisted). Unlike the 4WJ, which has 3 structural isomers, the 6WJ potentially has 40 structural isomers arising from one set of sequences. To learn more about the structural dynamics of these 6WJs, we built and simulated 14 unique isomers of the 6WJ comprising 7 unique motifs using all-atom molecular dynamics (MD). The first isomer of each individual motif was simulated, and all of the isomers for the **TO** and **PP** motifs were simulated to explore structural and energetic differences between isomers of the same motif.

From the MD analysis, we first quantified the coarse-grained geometry of each simulated 6WJ isomer by calculating stacked duplex-duplex angles, and determined that isomers of the same motif have similar duplex-duplex angles that are distinguishable from other motifs. Next, we compared the free energies of each simulated 6WJ isomer by using the MM-PBSA method, and we show that in isomers of the same motif, structural isomerization results in differing energetic favorability, likely due to the sequence-dependent effects based on the different duplex stacking partners.^71^ Similarly, we showed that different motifs have distinct energetics, where the twisted and planar parallel motifs are the most energetically favorable, due to the minimization of backbone-backbone interactions between duplexes, previously seen in isomers of the DX tile.^6^ The most favorable twisted motifs have duplex stacking angles similar to the 60*^◦^* anti-parallel 4WJ^13^ isomer, generally the most stable conformation of the 4WJ. In addition, the **TO1** isomer was the only isomer outside of the twisted and planar parallel configurations that had a relatively favorable free energy, and this is the motif that we use as a junction-level building block in the next section. Interestingly, the top orthogonal isomers have duplex stacking angles that are not completely orthogonal, preferring angles skewed slightly from 90*^◦^*, which is due to a mismatch between the minor and major grooves on the top and bottom duplexes of the stacked structure. Finally, from a base-pair step (BPS) parameter analysis, we have highlighted different important structural features on each 6WJ isomer that display how the DNA strands adjust into different stacked conformations and the implication of large deviations throughout the duplexes on each motifs favorability to form.

A major goal of quantifying the available conformations in a stacked 6WJ is to determine how to use these 6WJ isomers as junction-level building blocks for novel DNA nanostructures. For this work, we focus on the creation of a 2D square lattice using four **TO** motifs connected in a 2 by 2 tile arrangement. This 2D lattice allows for junction-level orthogonal directional changes in a DNA nanostructure at a resolution less than 10 nm, a smaller mesh size than seen in the literature for structures using the 4WJ and multi-dimensional vertices as a building block. ^77,78^ Experimental gel electrophoresis shows that this 2D lattice folds into a monodisperse population when using a single central staple strand to connect the four 6WJ motifs. Next, dynamic light scattering shows a single population of the 2D lattice at a size of 13.5 nm, very close to the predicted value of around 12 nm for each edge length. Lastly, atomic force imaging of the nanostructures confirms the correct size and shape of the tiles, and shows their slight deformation at the junction-level into structures skewed from orthogonality. An MD simulation of the 2D tile assembly confirms that the design prefers angles skewed from orthogonal, with interior angles closer to 100*^◦^* or 80*^◦^* than orthogonal, matching the images from AFM. We further note that this is the first 2D tile assembly built using an orthogonal topology at the junction-level for DNA nanotechnology, and this allows us to achieve much smaller mesh sizes than seen with the 4WJ motif.

This study lays the groundwork for the future quantification of the isolated 6WJ, with computational studies likely focusing on the conformational landscape using exploratory methods such as REMD^74^ and thermodynamic integration.^75^ In addition, we propose to design and build additional 1D, 2D, and 3D nanostructures that can be tiled from the individual 6WJ motifs, as we focused here only on the potential for the top orthogonal motif to tile into a 2D square lattice. From these initial studies we hope to introduce the 6WJ as a building block to the forefront of DNA nanotechnology and accelerate its use into new and exciting DNA structures.

## Computational Methods

### DNA six-way junction atomic models

Each atomic model was constructed in Biovia Discovery Studio,^79^ by starting from a 6-armed open star we call the **Open** motif, analogous to the 4WJ open motif. From this **Open** structure, we identified all stacked 6WJ motifs consisting of three connected duplex arms by rotating and translating the duplexes in the initial structure. These include the distinct planar parallel (**PP**), top orthogonal (**TO**), bottom orthogonal (**BO**), right-handed parallel (**RP**), left-handed parallel (**LP**), right-handed twisted (**RT**), and left-handed twisted (**LT**) motifs which are discussed in detail in the *Results and Discussion* section. Different isomers of each motif were created by modifying the sequence of each original motif. The square lattice atomic model was created by combining atomic models of four separate TO motifs.

### All-atom molecular dynamics simulations

All-atom MD was chosen to simulate the dynamics of our 6WJ motifs, as we are more interested in the equilibrium dynamics of each isomer than the longer time scale biological dynamics. Each structure was loaded into the AMBER 20^67^ program tleap, implicitly solvated with the pairwise Generalized Born model.^80–82^ The DNA.OL15^83^ force field was applied and each structure was minimized with a cutoff of 20Å over 1000 steps. The minimized structures were then neutralized with Mg^2+^ ions and solvated in a 20Å box of TIP3P water,^84^ where 12.5 mM of MgCl_2_ were added randomly to the system. The solvent and ions were then minimized for a total of 5000 steps, while the solute was held in place with a restraint of 500 kcal/(mol·Å). Next, the entire solvated system was minimized for a total of 5000 steps. The structures were then heated from 0 to 300 K and equilibrated over 50000 time-steps at 2 fs for a total of 100 ps. During heating, constant volume periodic boundaries were applied, as well as a weak force constant on the DNA of 10 kcal/(mol·Å) and a cutoff of 12Å. Temperature regulation via Langevin dynamics^85^ was used with a collision frequency of 5 ps*^−^*^1^ with the SHAKE algorithm^86^ applied. After heating the system was allowed to equilibriate for 400 ps with a constant temperature of 300K, with constant pressure periodic boundaries using a Berendsen barostat^87^ at a pressure of 1 atm and a relaxation time of 2.0 ps. Temperature regulation via Langevin dynamics^85^ was used, along with the same collision frequency, cutoff, and SHAKE settings used in heating. After heating and equilibration, three replicates of each structure were run for 150 ns with randomized input velocities. Coordinates were written out every 1000 steps or 2 ps. After completing the MD runs, the CPPTRAJ^88^ program was used to combine the trajectory file for each duplicate and strip the files of the solvent. The first 25 ns of simulation time was cut from each replicate to ensure an equilibrium state had been reached. A total of 1250 frames from each replicate or 3750 frames per structure were kept, which is one frame per 100 ps. The RMSD for each replicate was also calculated using CPPTRAJ,^88^ extracted and graphed using Python’s matplotlib.^89^ For the square lattice MD simulation, only one replicate of 150 ns was performed, but all other parameters were identical to the above.

### Structural analysis of DNA six-way junctions

The Python package ProDy^90,91^ was used to analyze structural properties for each replicate simulated using all-atom MD. For each frame analyzed, the axis along each DNA duplex 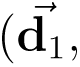 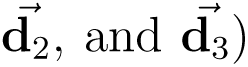 was calculated with C = UΛ U *^T^*, where C is the covariance matrix describing the atomic coordinates of all the atoms in the separate duplexes, Λ is the the diagonal matrix storing the eigenvalues of C, and U is a set of eigenvectors of C. The eigenvector corresponding to the largest eigenvalue is assumed to be the orientational vector following the duplex axis 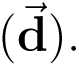 Finally, the orientation of each vector was made to match the direction of the 5’ to 3’ end of the scaffolding strand by comparison with a reference vector calculated by extracting coordinates of specific bases. The duplex-duplex angles for all duplex combinations in each 6WJ motif were then calculated by taking the following dot products: 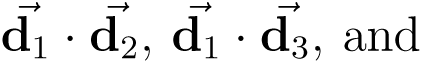 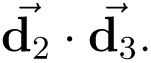

### Free energy analysis of DNA six-way junctions using MMPBSA

Free energy calculations were conducted using the Molecular Mechanics Poisson-Boltzmann Surface Area (MM-PBSA) method^68^ as implemented in AMBER 20.^67^ The MM-PBSA method estimates the Gibbs free energy of solvated molecules as G*_solvated_* = E*_gas_* +G*_solvation_* − T S*_solute_*, where E*_gas_* is the gas-phase contribution of the solute to free energy, G*_solvation_* is the free energy change upon solvation, T is the temperature, and S*_solute_* is the solute entropy. As calculating the energetics of an explicitly solvated water box would be too computationally expensive, the MM-PBSA method implicitly solvates the DNA molecule in a continuum solvant^92^ using the solvent-accessible surface area. The energetic contribution of the solvation free energies are broken into polar and nonpolar contributions as G*_solvation_* = G*_polar_* + G*_nonpolar_*. The nonpolar solvation effects are modeled as G*_nonpolar_* = γSA + b, where SA is the surface area of the solute and γ and b are constants.^92^ The polar contributions can be calculated from multiple methods but MM-PBSA specifically employs the PoissonBoltzmann equation as ∇ɛ(r)∇ϕ(r) + 4πρ(r) = 0, where *ɛ*(r) is the dielectric function, ϕ(r) is the electrostatic potential, and ρ(r) is the charge density caused by the solute.^92^ The Poisson-Boltzmann equation was solved numerically over 1000 iterations per frame, at an ionic strength of 157.19 mM, with a 0.5 Å finite difference grid, 1.4 Å solvent probe radius, and a dielectric constant of 1.0 and 80.0 for the solute and solvent, respectively.

Following this, entropy corrections were performed to more accurately calculate the free energy of each 6WJ motif. The entropy was estimated through normal mode calculations which were conducted on 750 evenly sampled frames per motif. The normal mode calculations consist of solvation of each frame using the Generalized-Born model^81,82^ with dielectric constants of 1.0 and an ionic strength of 157.19 mM. Rigid rotor models were used to find translation and rotational entropies of the solute. For vibrational entropies each frame is then minimized over 5000 cycles with a convergence criteria of.01 kcal/mol, and a Hessian matrix was diagonalized to find the vibrational frequencies. The vibrational entropy can be estimated with standard vibrational partition functions, after which the total summed entropy is subtracted from G*_solvated_*.

### DNA six-way junction base-pair step analysis

The X3DNA^93,94^ program was used to extract the widely used 3DNA base-pair step parameters over each combined 6WJ motif trajectory. For each frame a set of “idealized” base coordinates are fit to the trajectory using a least-square regression procedure,^95^ with a 3 x 3 matrix describing the x, y, and z axis of the base. Axes describing base-pairs can be derived from a rotation and averaging of the individual base axes.^96^ Likewise, base-pair step (BPS) axes can be calculated through a rotation and average of each base-pair axes. The x-axis of a base-pair points towards the major groove, with the y-axis parallel to a line between the C1’ carbons on the sugars. The z-axis is then defined as the cross product of x and y. For base-pair steps the x-axis still points towards the major groove, with the y-axis the average between the two individual base-pair y-axes, and the z-axis is defined as the cross product of x and y.^97^ From the axis definitions, translational parameters (rise, shift, and slide) and rotational parameters (tilt, roll, and twist) can be calculated for each base-pair step in all frames. After calculating and averaging these parameters for each BPS they were subtracted from a reference value^98^ corresponding to the average B-form DNA duplex structure for each BPS, and these base-pair step deviations are reported.

## Experimental Methods

### Oligonucleotides

All modified (fluorescently labeled) or non-modified oligonucleotides used in this study to assemble the various nanostructures were acquired from Integrated DNA Technologies (IDT) and used without further purification.

### Sequence design

The sequence used to assemble the DNA 6-way junction designed in this study were designed using the Tiamat 2 software. ^55^ The sequences were designed to be orthogonal to each other to avoid any mispairing. The sequences were further validated using NCBI BLAST.^64^ All the sequences designed and used in this study are presented in Tables S2 to S4.

### DNA motif assembly and characterization

To fold the different motifs, multiple strands (modified or non-modified) were mixed at equimolar concentration (5 micromolar) in Tris-Acetate EDTA-MgCl buffer (40 mM Tris, 20 mM acetic acid, 2 mM EDTA, 12 mM MgCl_2_, pH 8.0) to prepare a stock solution for each variant tested. The oligonucleotide mix was then slowly annealed from 95°C to 4°C overnight using a BioRad T100 thermocycler. The folded structures were then stored at 4°C prior further use. To confirm proper folding of the structures and the absence of bi-products, the annealed structures were characterized via agarose gel electrophoresis with 1.5% agarose gels preloaded with ethidium bromide and images were taken using an Azure Gel Imager C150 (Azure Biosystems).

### Fluorescence resonance energy transfer (FRET) assay

Two of the oligonucleotides used for the assembly of the non-modified DNA 6WJ motifs were modified with the FRET donor dye Fluorescein (6WJ-1 and 6WJ-5, Table S3) and 3 oligonucleotides were modified with the FRET acceptor dye TAMRA (6WJ-2, 6WJ-3, and 6WJ-6, Table S3) to create multiple configuration of the FRET pairs. We tested 6 different FRET pairs (3 with 6WJ-1 as donor and 3 with 6WJ-5 as donor). After folding, the different variants were diluted in PBS and fluorescence measurements were performed in a Tecan Safire2 fluorescent plate reader with an excitation wavelength set at 455 nm (FAM excitation) and the emission collected between 505 nm and 700 nm (FAM and TAMRA emission). The FRET efficiency and the percentage of intact structures was calculated using the equation E = 1 − I*_DA_*/I*_D_*, where E is the FRET efficiency, I*_DA_* is the intensity of the donor emission in presence of both donor and acceptor, and I*_D_* is the intensity of the donor in absence of the acceptor.

### Dynamic Light Scattering (DLS)

Dynamic light scattering (DLS) experiments were performed with a NanoZetaSizer (Malvern Instruments, Ltd) to measure the hydrodynamic diameter of the tile assembled. Technical replicates (n = 3) of 250 nM tile were performed in folding buffer at 25°C.

### Atomic force microscopy (AFM)

The tiles were imaged using a Bruker Dimension Icon + ScanAsyst AFM instrument. A mica surface (Electron Microscopy Sciences) was freshly cleaved and 40 µL of 5-10 nM DNA-NPs in FB were added to the surface. Following a 10 min incubation, fresh folding buffer was added to gently wash the surface several times. AFM images were then captured in folding buffer supplemented with 10 mM NiCl_2_ as imaging buffer, ScanAsyst mode, and a SNL-10 probe (Bruker). Images were processed using Gwyddion.^99^ In some instances (indicated in the caption), the surface was treated with 0.1 mg/ml Poly-D-Lysine (Thermo Fisher Scientific) and rinsed with ultrapure water and dried with compressed air prior to load the DNA-NPs on the surface.

## Acknowledgement

We thank the National Science Foundation under NSF-CISE award #2312215 for partially supporting this work. We thank the UNM Center for Advanced Research Computing, supported in part by the National Science Foundation, for providing the high performance computing and large-scale storage resources used in this work.

