## Supplementary Information for "DNA six-way junction conformations and their use in a 2D square lattice"

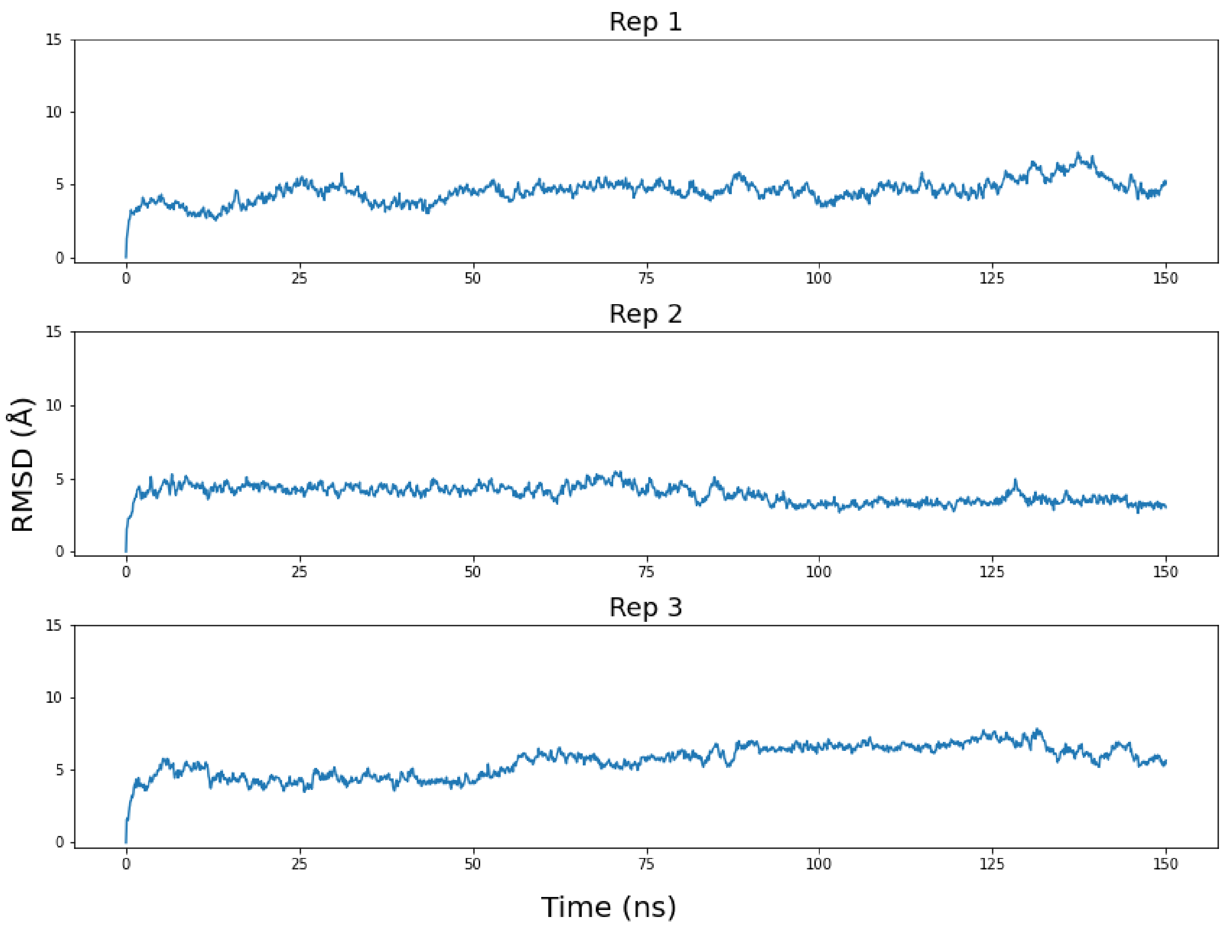


**Figure S1: RMSD of the TO1 isomer over 150 ns.** The RMSD of each Top Orthogonal Isomer 1 replicate over its 150 ns trajectory.

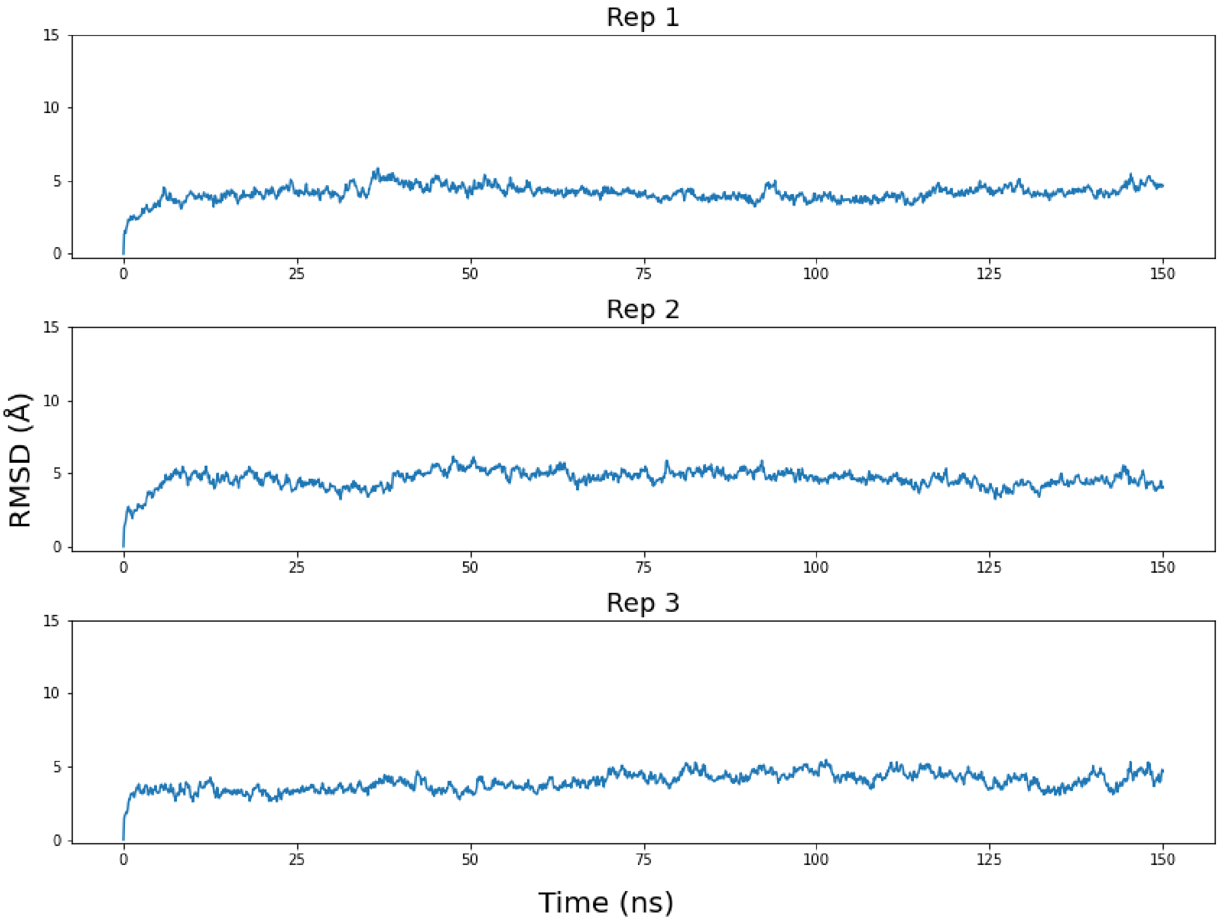


**Figure S2: RMSD of the TO2 isomer over 150 ns.** The RMSD of each Top Orthogonal Isomer 2 replicate over its 150 ns trajectory.

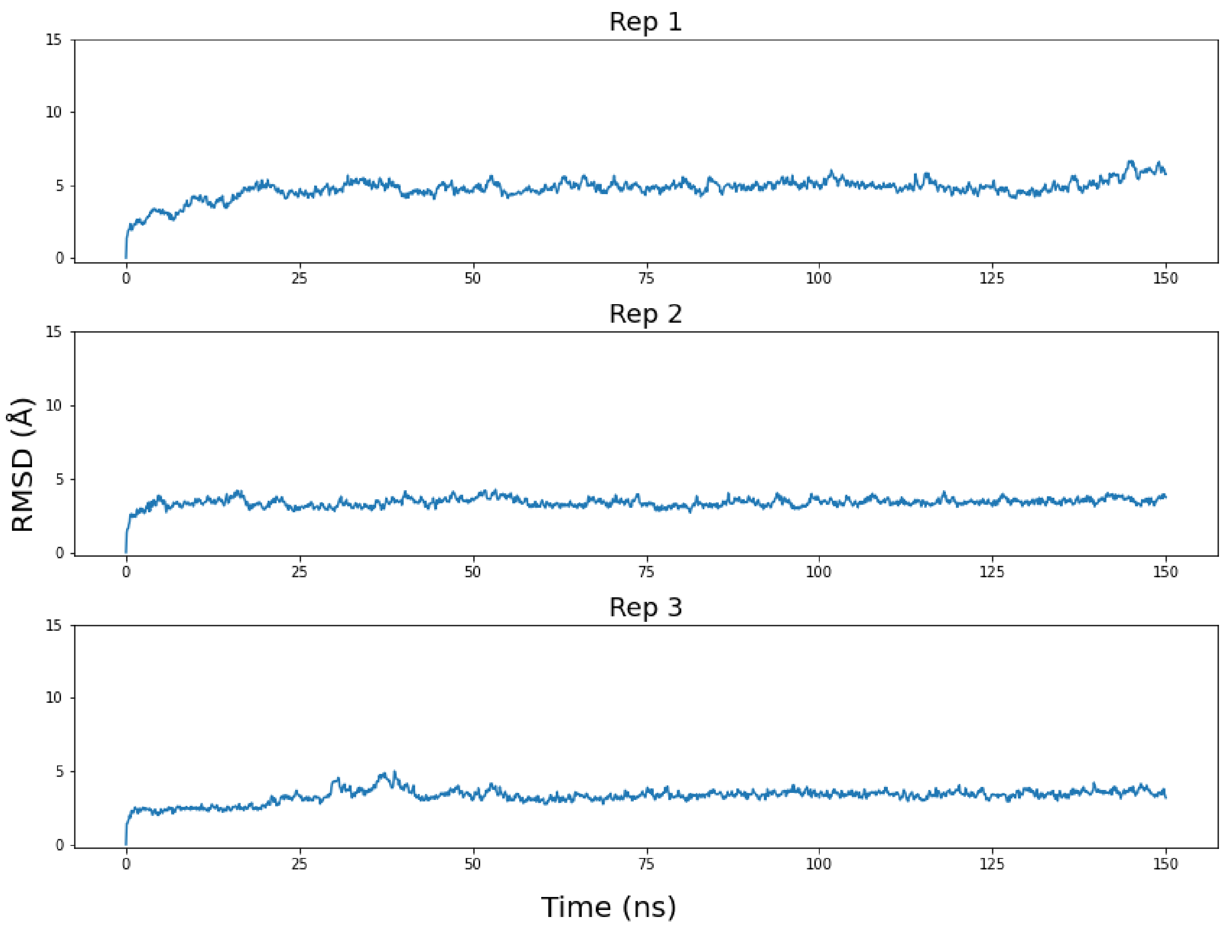


**Figure S3: RMSD of the TO3 isomer over 150 ns.** The RMSD of each Top Orthogonal Isomer 3 replicate over its 150 ns trajectory.

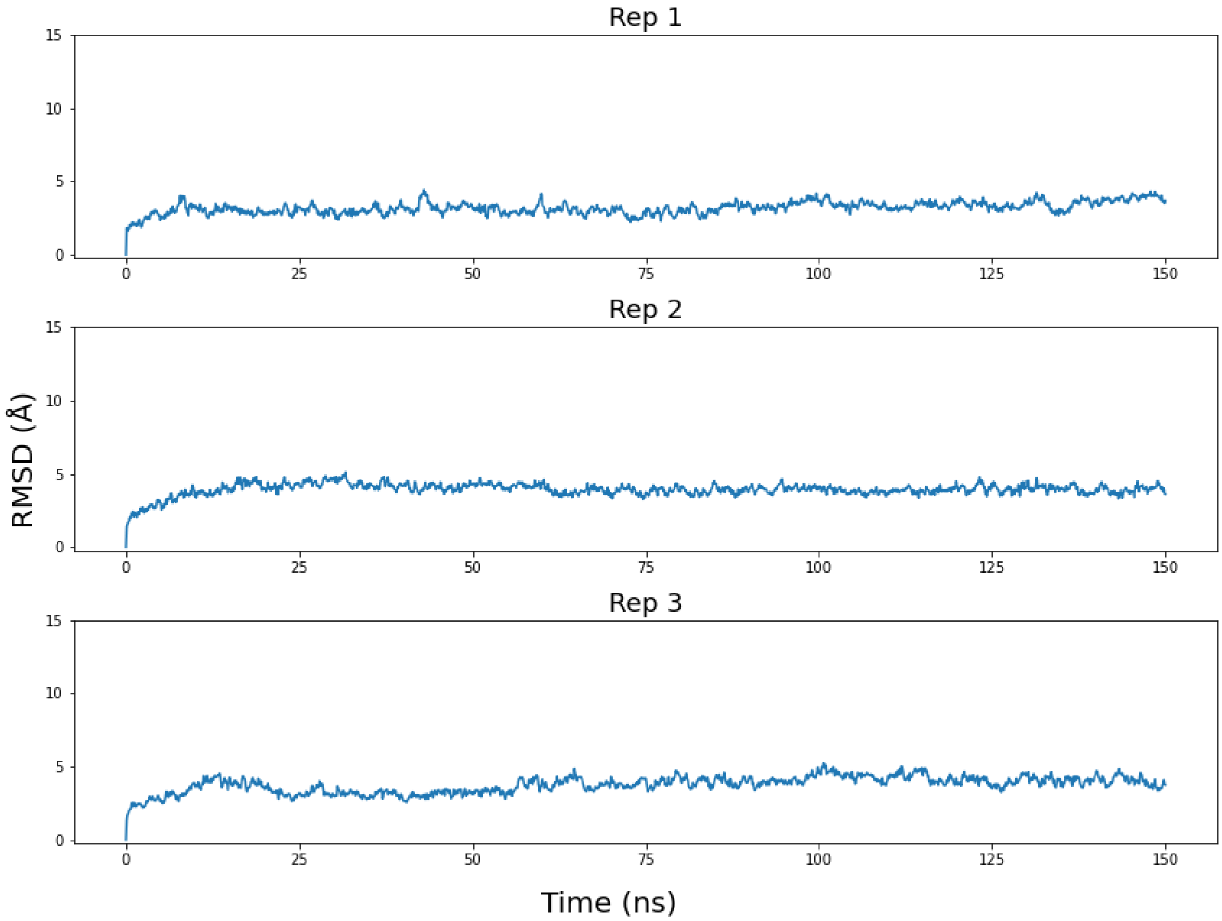


**Figure S4: RMSD of the TO4 isomer over 150 ns.** The RMSD of each Top Orthogonal Isomer 4 replicate over its 150 ns trajectory.

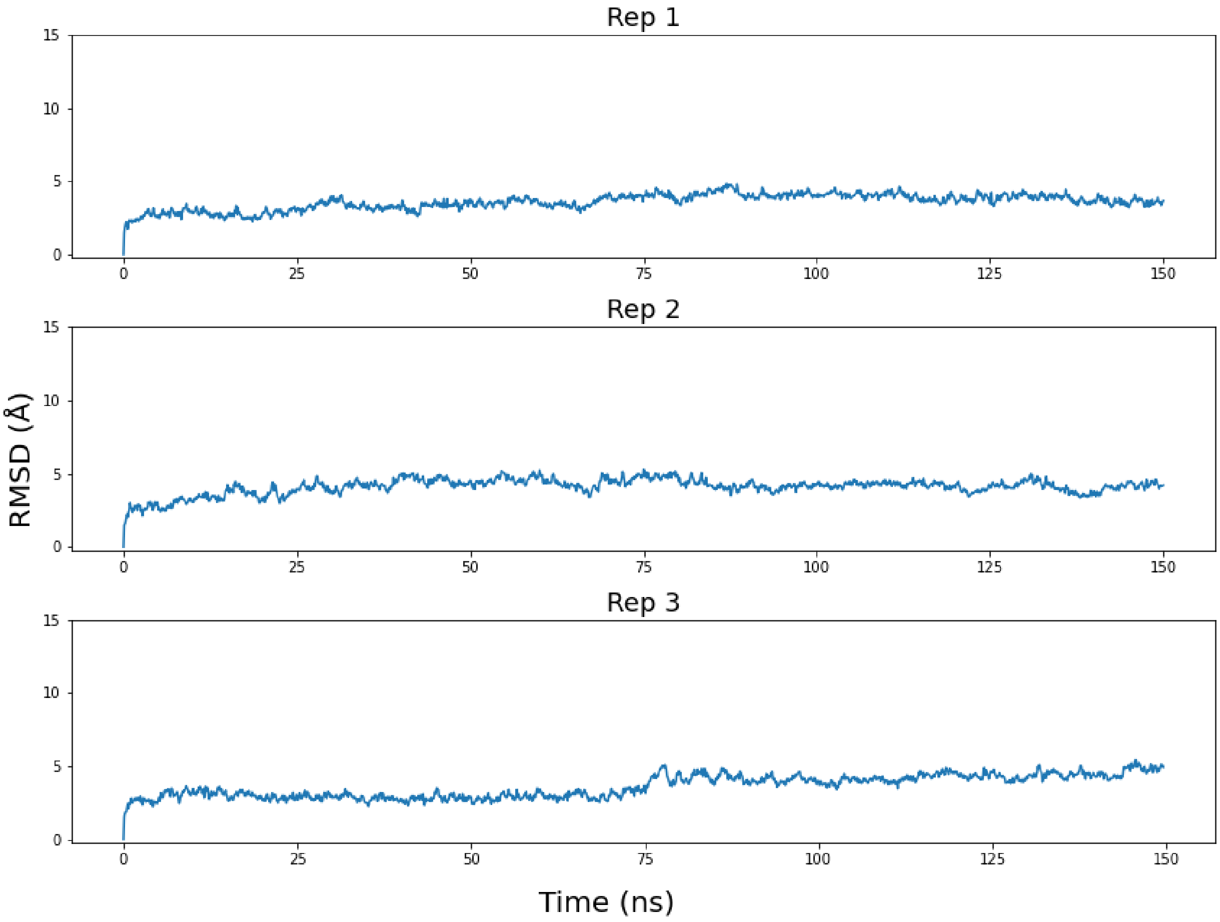


**Figure S5: RMSD of the TO5 isomer over 150 ns.** The RMSD of each Top Orthogonal Isomer 5 replicate over its 150 ns trajectory.

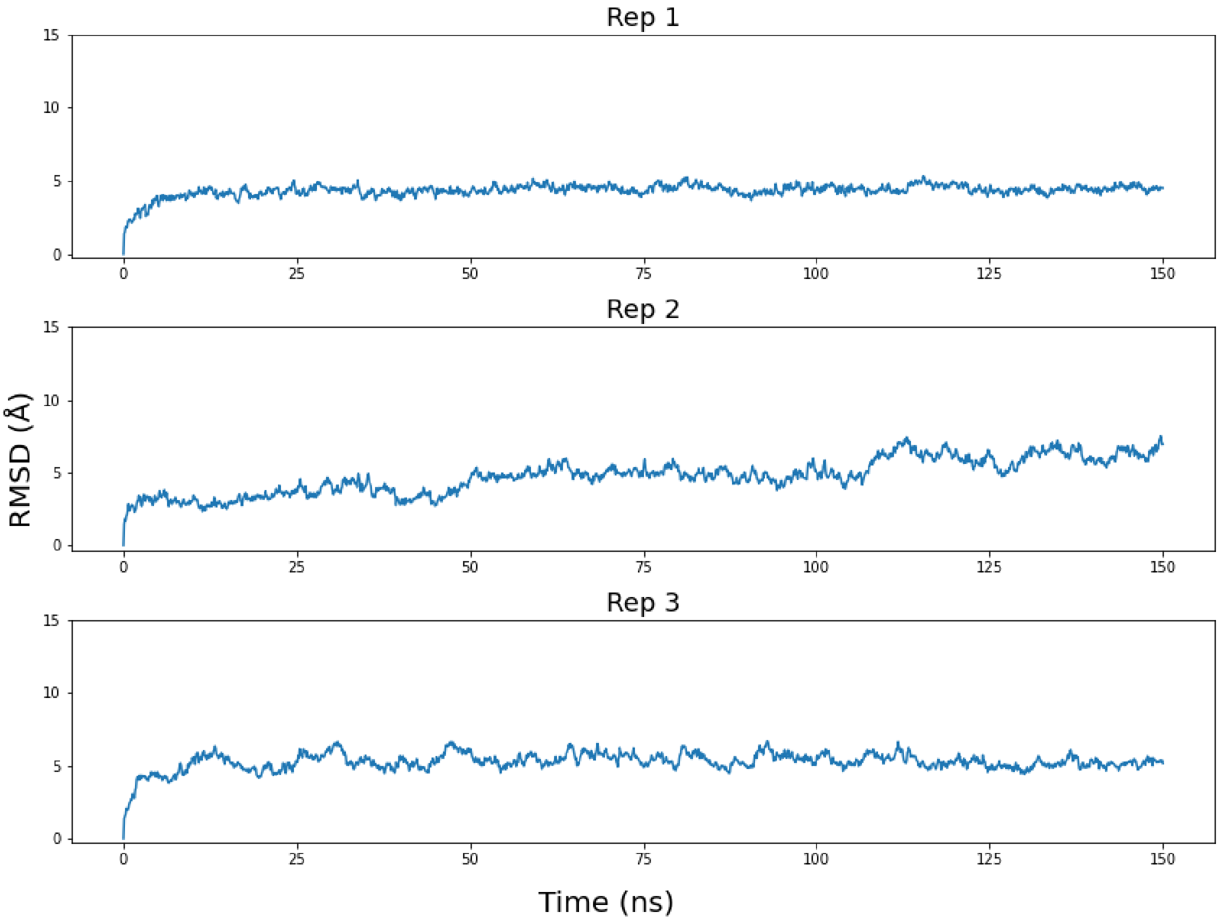


**Figure S6: RMSD of the TO6 isomer over 150 ns.** The RMSD of each Top Orthogonal Isomer 6 replicate over its 150 ns trajectory.

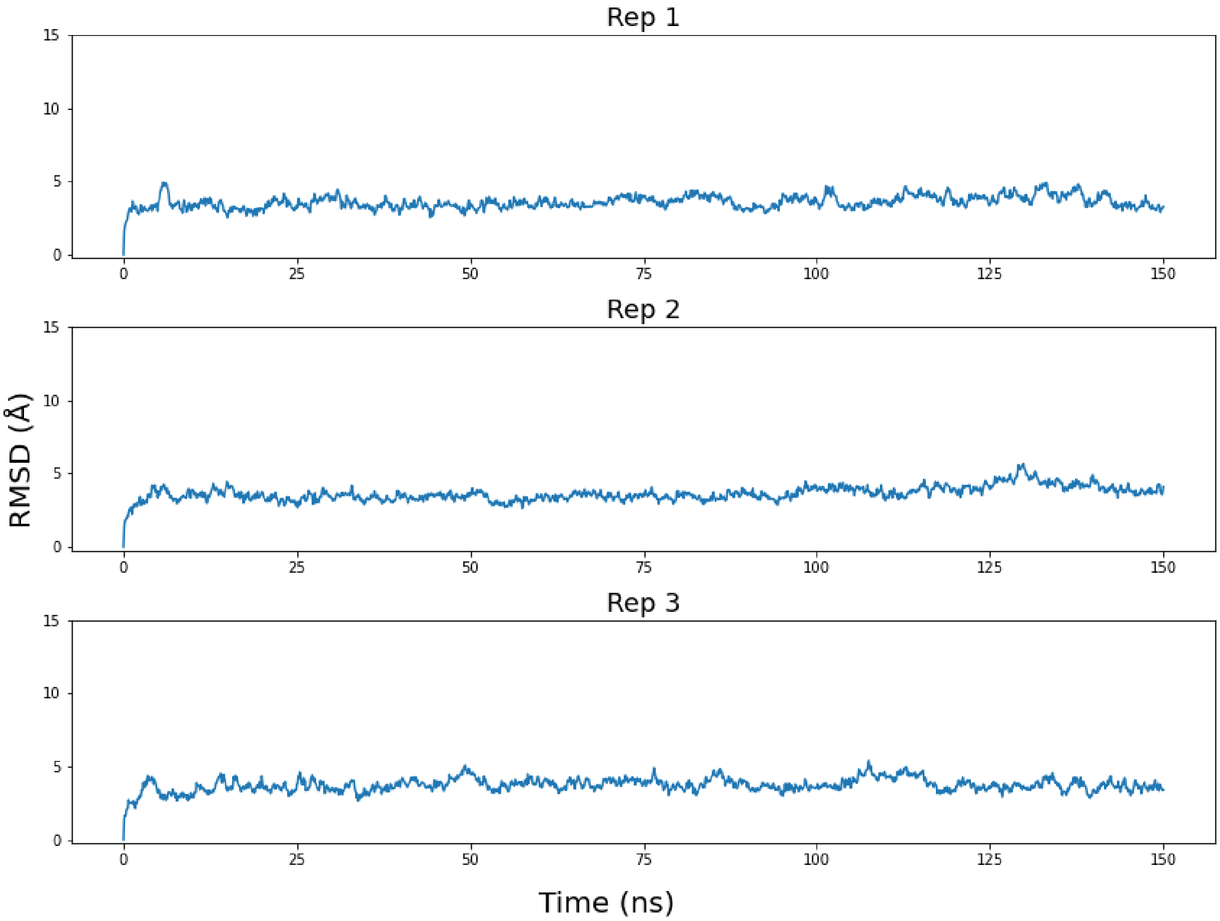


**Figure S7: RMSD of the BO3 isomer over 150 ns.** The RMSD of each Bottom Orthogonal Isomer 3 replicate over its 150 ns trajectory.

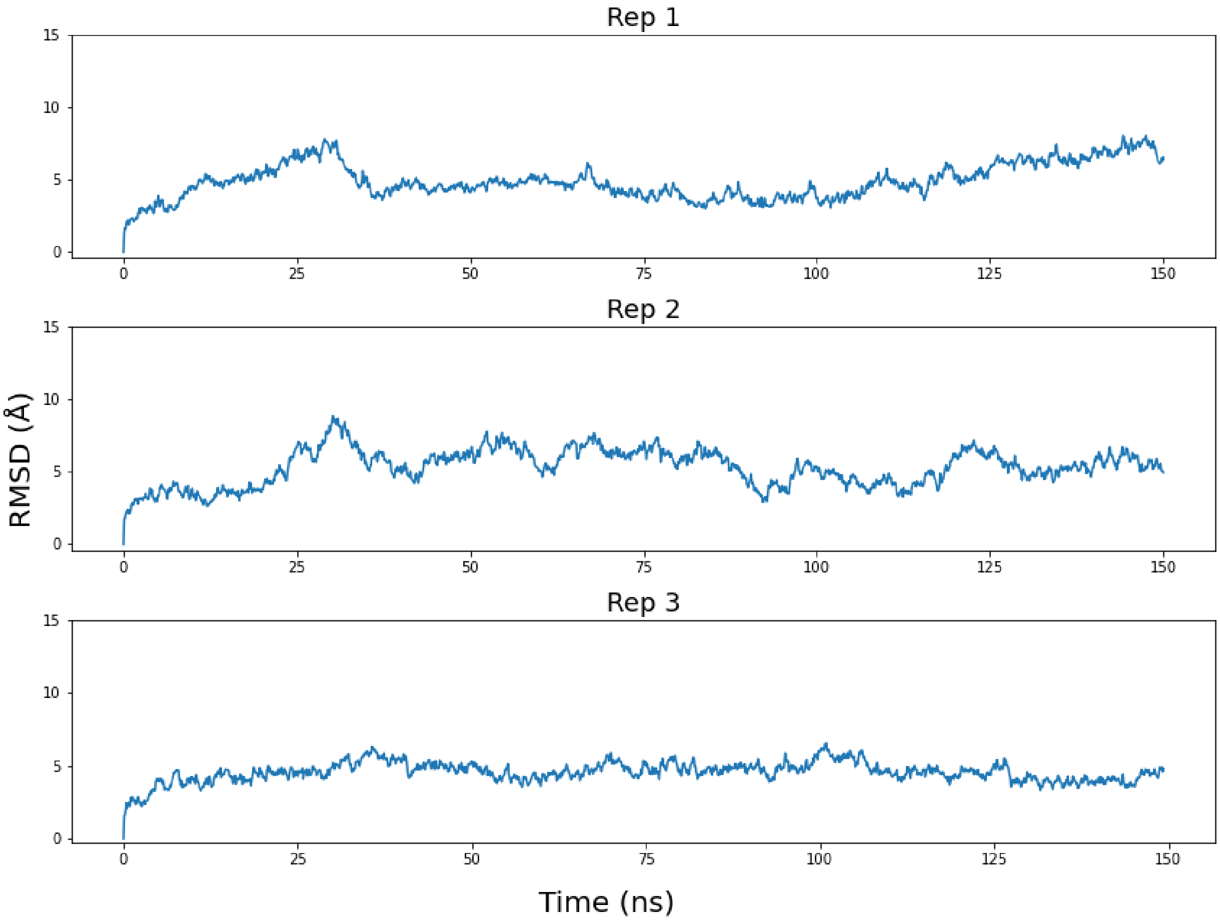


**Figure S8: RMSD of the PP1 isomer over 150 ns.** The RMSD of each Planar Parallel Isomer 1 replicate over its 150 ns trajectory.

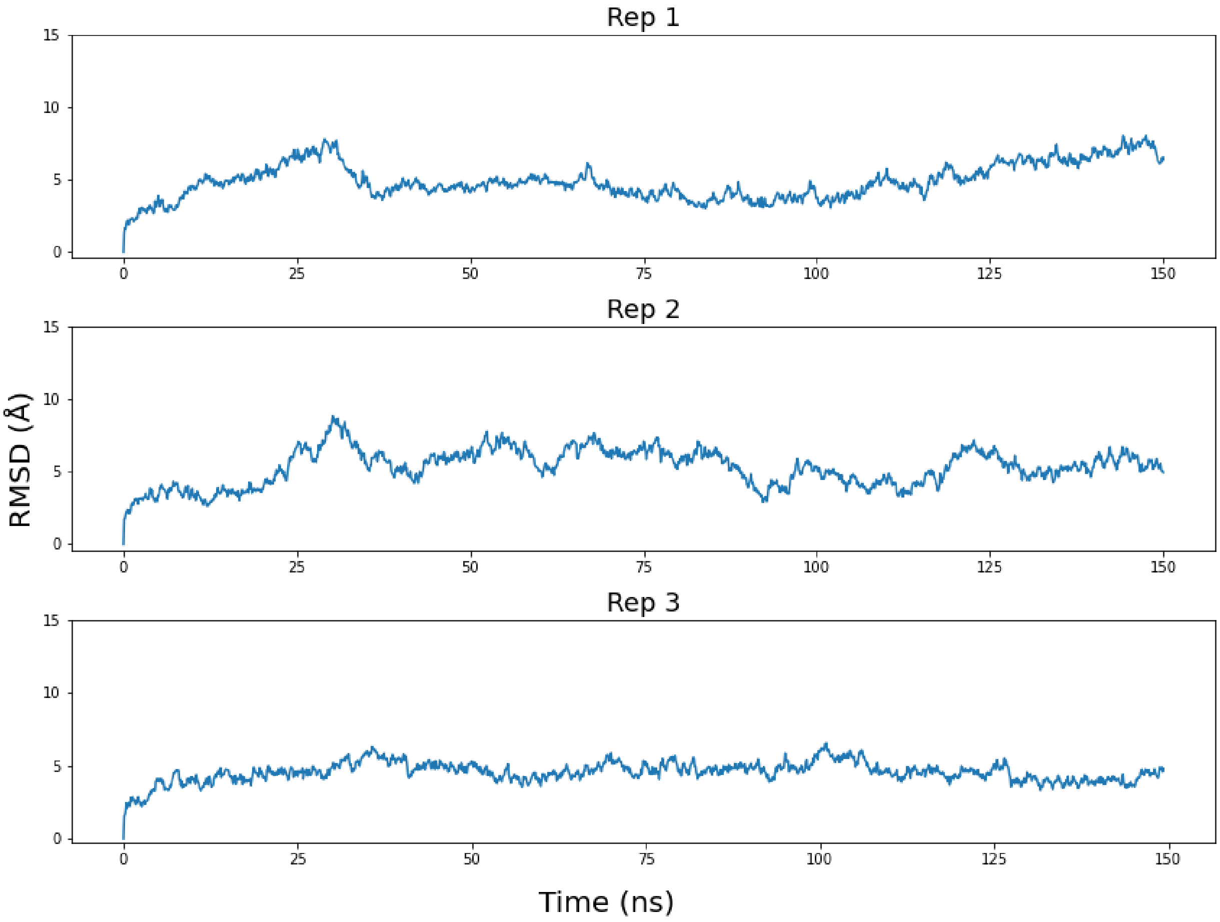


**Figure S9: RMSD of the PP2 isomer over 150 ns.** The RMSD of each Planar Parallel Isomer 2 replicate over its 150 ns trajectory.

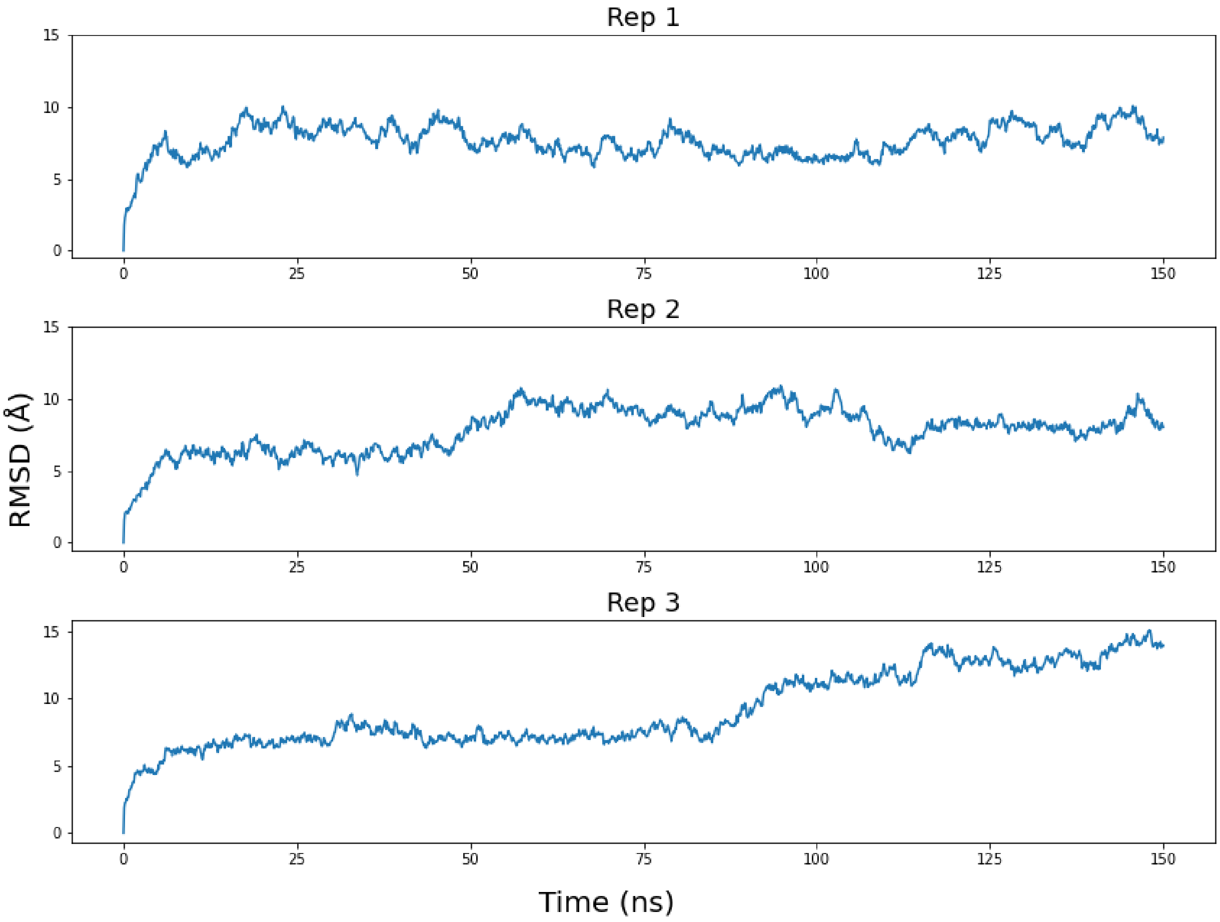


**Figure S10: RMSD of the PP3 isomer over 150 ns.** The RMSD of each Planar Parallel Isomer 3 replicate over its 150 ns trajectory.

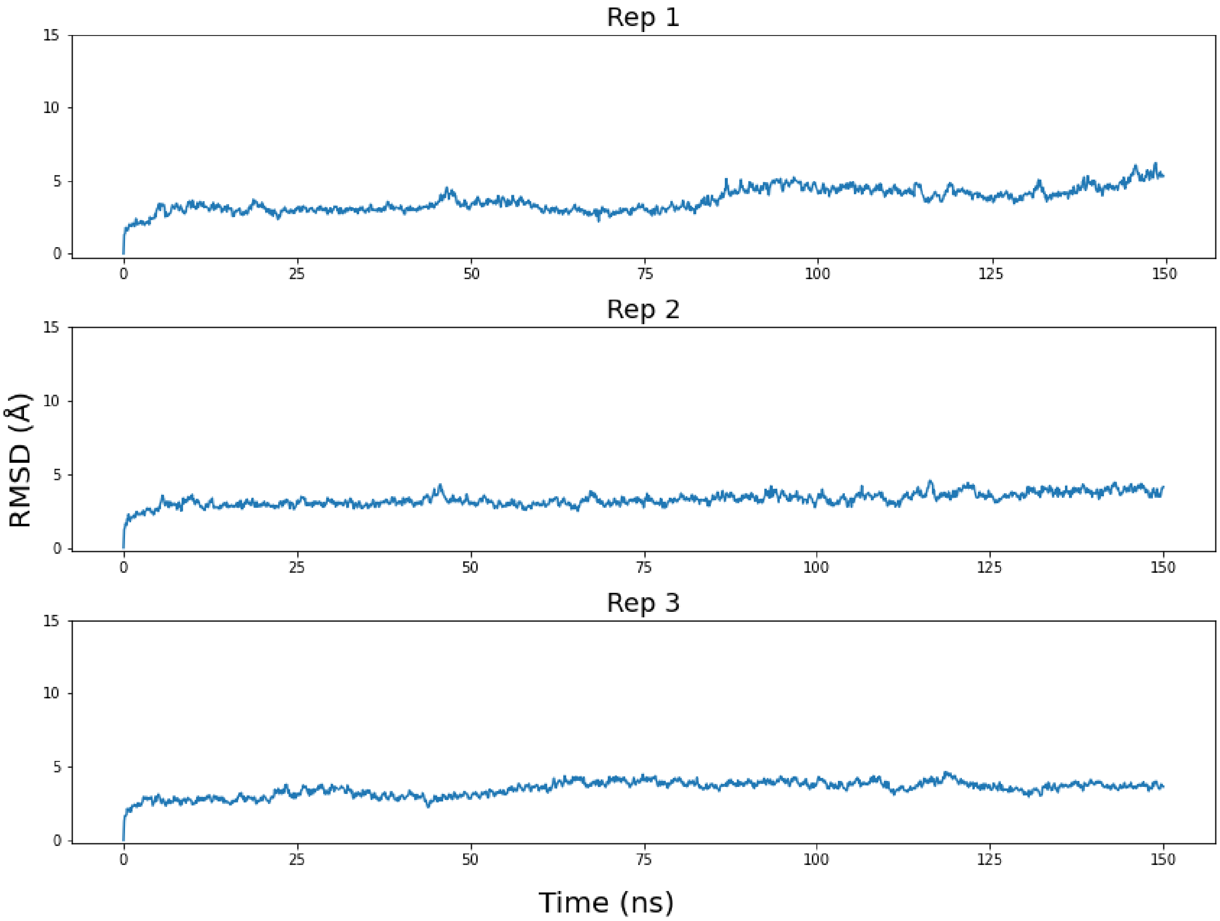


**Figure S11: RMSD of the RP1 isomer over 150 ns.** The RMSD of each Right-Handed Parallel Isomer 1 replicate over its 150 ns trajectory.

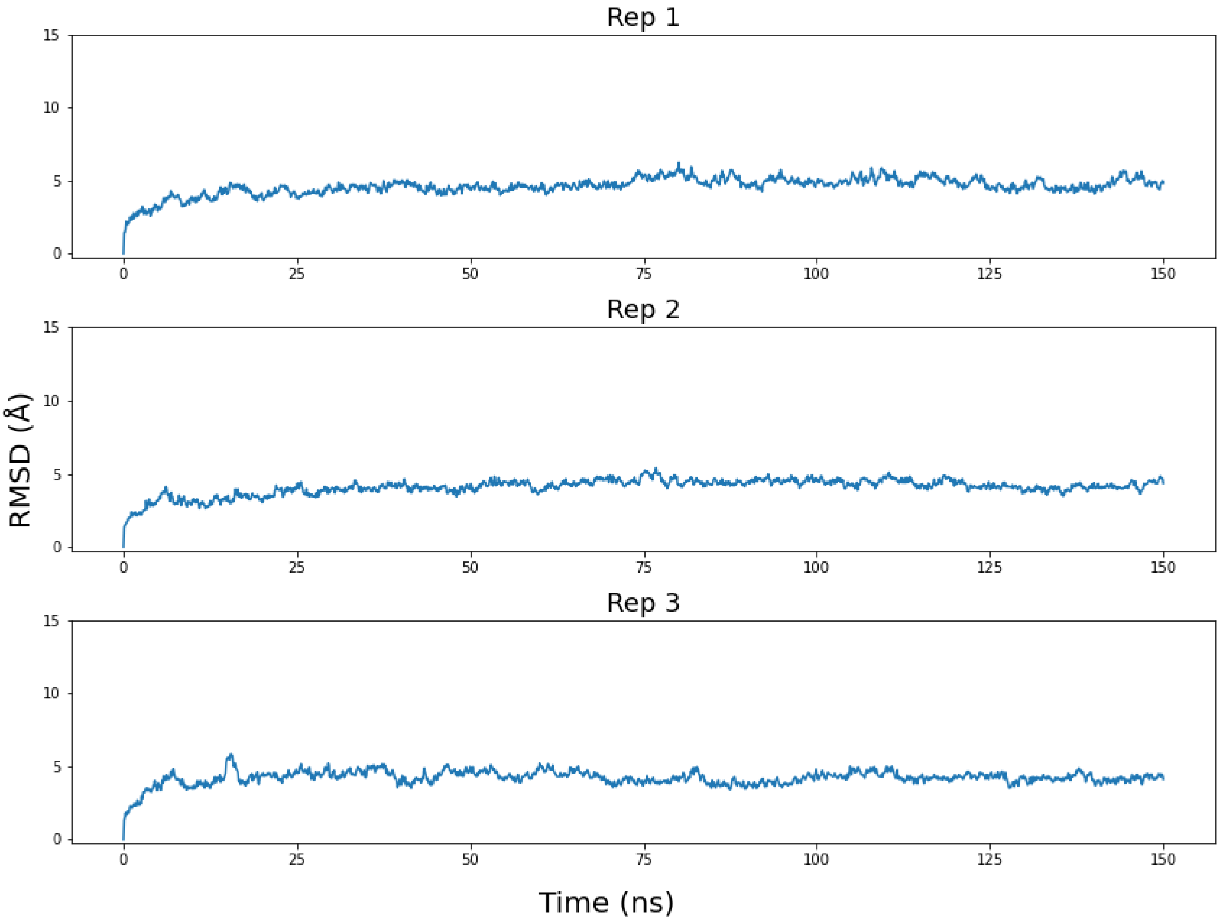


**Figure S12: RMSD of the LP1 isomer over 150 ns.** The RMSD of each Left-Handed Parallel Isomer 1 replicate over its 150 ns trajectory.

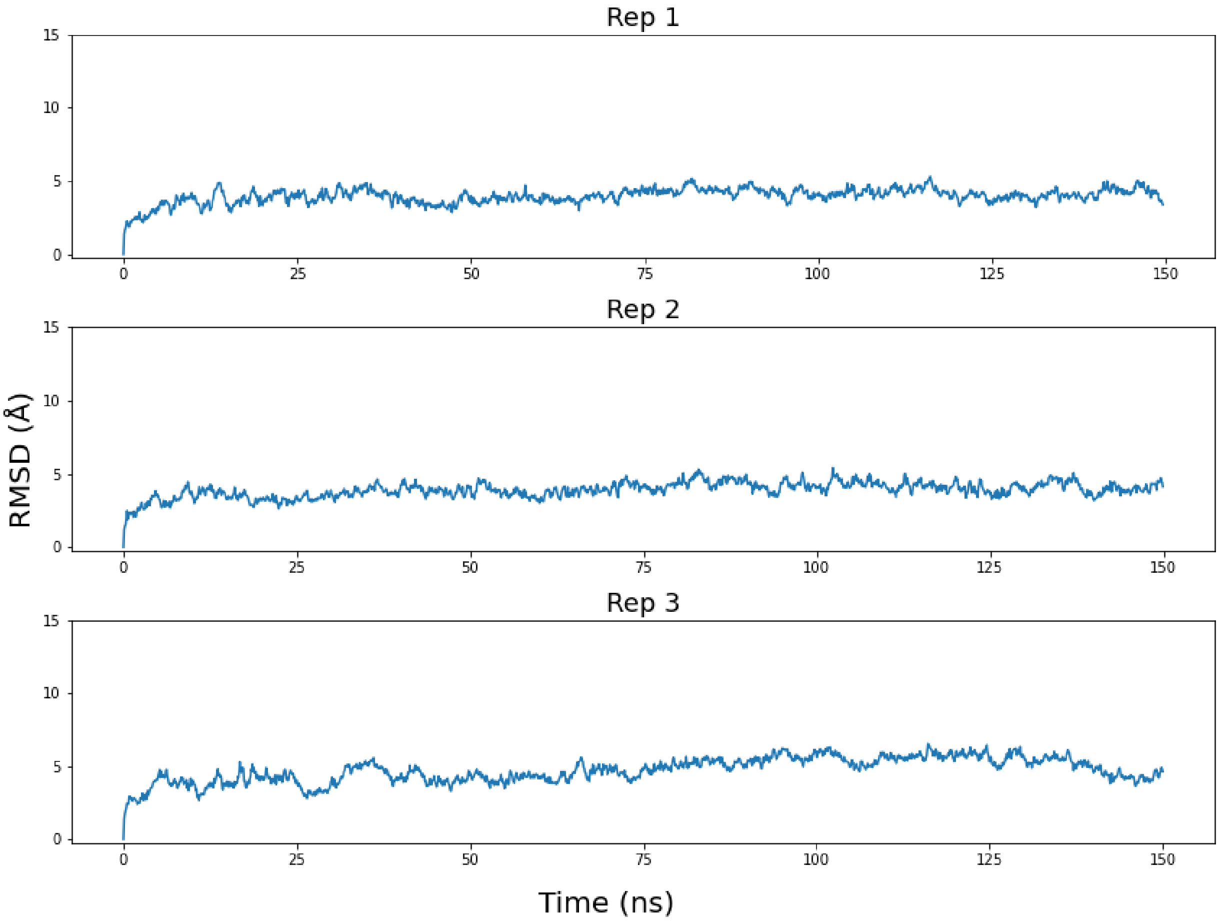


**Figure S13: RMSD of the RT1 isomer over 150 ns.** The RMSD of each Right-Handed Twisted Isomer 1 replicate over its 150 ns trajectory.

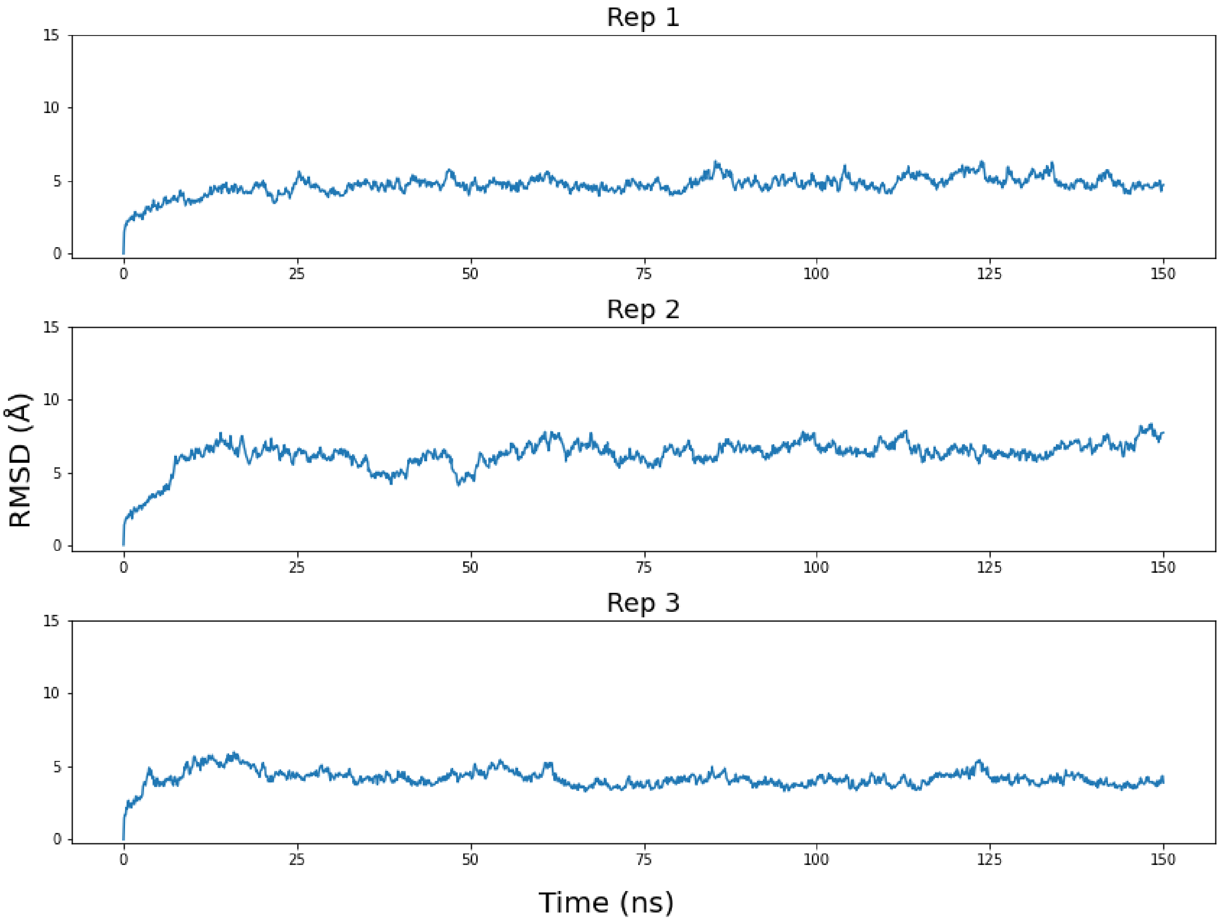


**Figure S14: RMSD of the LT1 isomer over 150 ns.** The RMSD of each Left-Handed Twisted Isomer 1 replicate over its entire 150 ns trajectory.

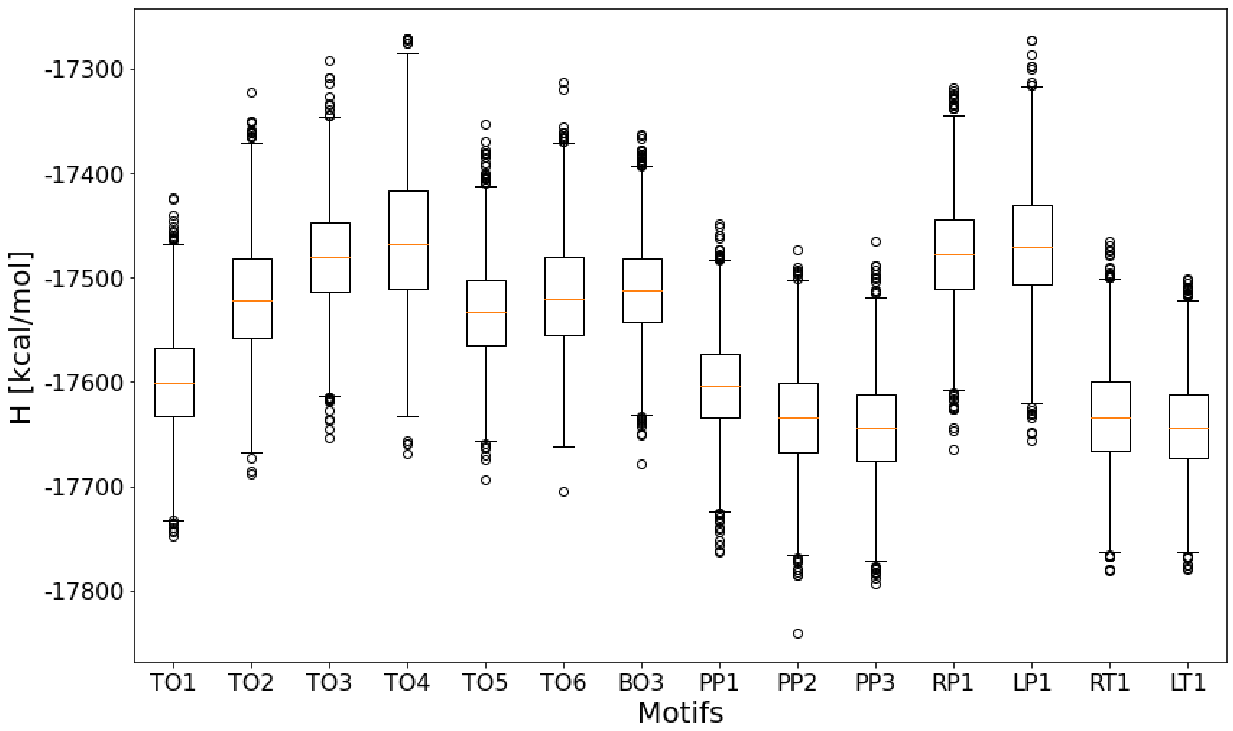


**Figure S15: $H$ values of simulated 6WJs.** A box-plot of the calculated $H$ values for all simulated isomers. The whiskers of each plot represent the minimum and maximum values excluding outliers, while the top and bottom edges of the box contain data points in the 25th-75th percentile with the orange line indicating the median. Outliers are determined to be greater than 1.5 multiplied by the inter-quartile range away from the top or bottom edges of the box.

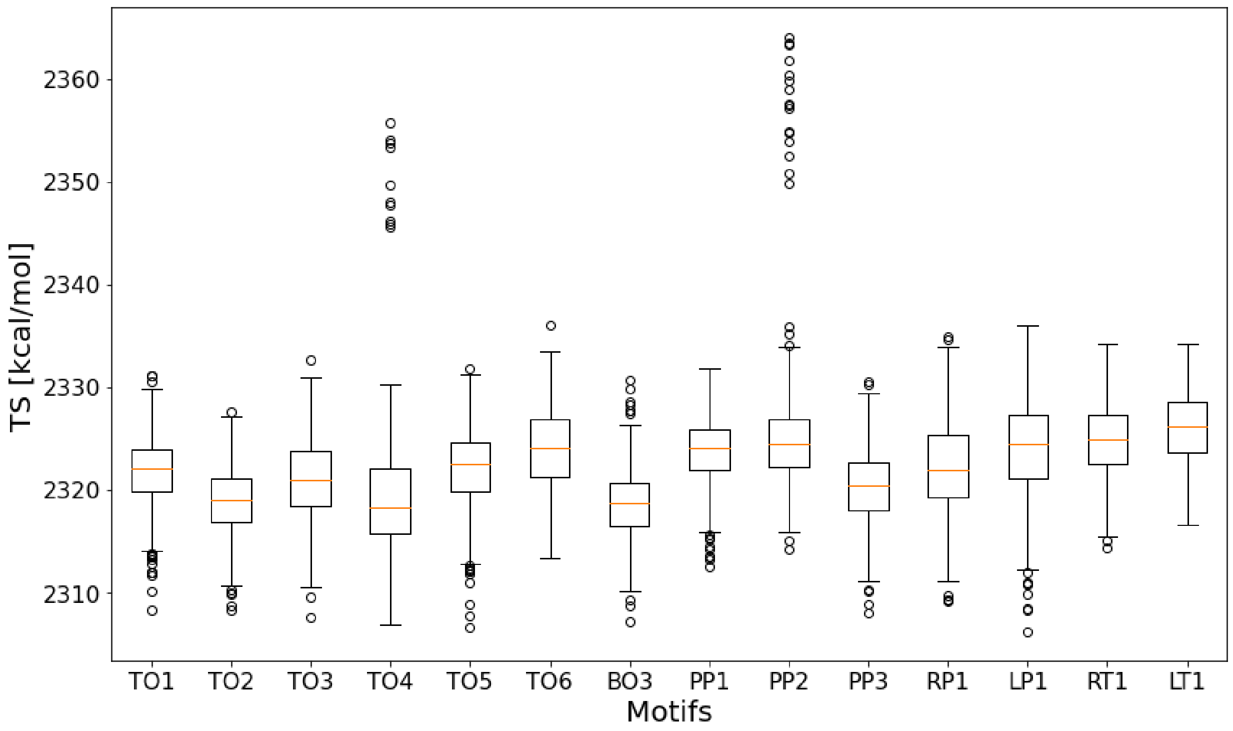


**Figure S16:** $TS$ **values of simulated 6WJs.** A box-plot of the calculated $TS$ values for all simulated isomers.The whiskers of each plot represent the minimum and maximum values excluding outliers, while the top and bottom edges of the box contain data points in the 25th-75th percentile with the orange line indicating the median. Outliers are determined to be greater than 1.5 multiplied by the inter-quartile range away from the top or bottom edges of the box.

**Table S1: Components of free energies of each 6WJ isomer.** The mean free energies broken down into the mean enthalpy and mean entropy terms each with 2 standard deviations noted in parentheses. All terms are in kcal/mol.

| Motif | $G$ | $H$ | $TS$ |
|---|---|---|---|
| TO1 | −20023.01(±97.21) | −17599.12(±96.36) | 2423.89(±12.83) |
| TO2 | −19936.87(±111.81) | −17518.99(±111.10) | 2417.88(±12.56) |
| TO3 | −19902.32(±101.14) | −17480.13(±99.92) | 2422.19(±15.67) |
| TO4 | −19881.19(±132.03) | −17462.79(±130.10) | 2418.40(±22.49) |
| TO5 | −19957.19(±93.31) | −17532.27(±92.11) | 2424.23(±14.88) |
| TO6 | −19945.46(±106.99) | −17517.44(±105.83) | 2428.02(±15.73) |
| BO3 | −19928.85(±89.39) | −17511.55(±88.43) | 2417.30(±13.07) |
| PP1 | −20031.17(±91.89) | −17603.43(±91.03) | 2427.74(±12.53) |
| PP2 | −20063.90(±98.83) | −17633.55(±95.89) | 2430.36(±23.93) |
| PP3 | −20064.56(±94.94) | −17643.90(±93.81) | 2420.65(±14.55) |
| RP1 | −19902.31(±99.51) | −17477.76(±97.99) | 2424.55(±17.34) |
| LP1 | −19897.21(±112.66) | −17468.98(±111.16) | 2428.23(±18.33) |
| RT1 | −20062.29(±97.90) | −17632.43(±96.85) | 2429.85(±14.3) |
| LT1 | −20074.41(±92.28) | −17642.36(±91.26) | 2432.04(±13.72) |

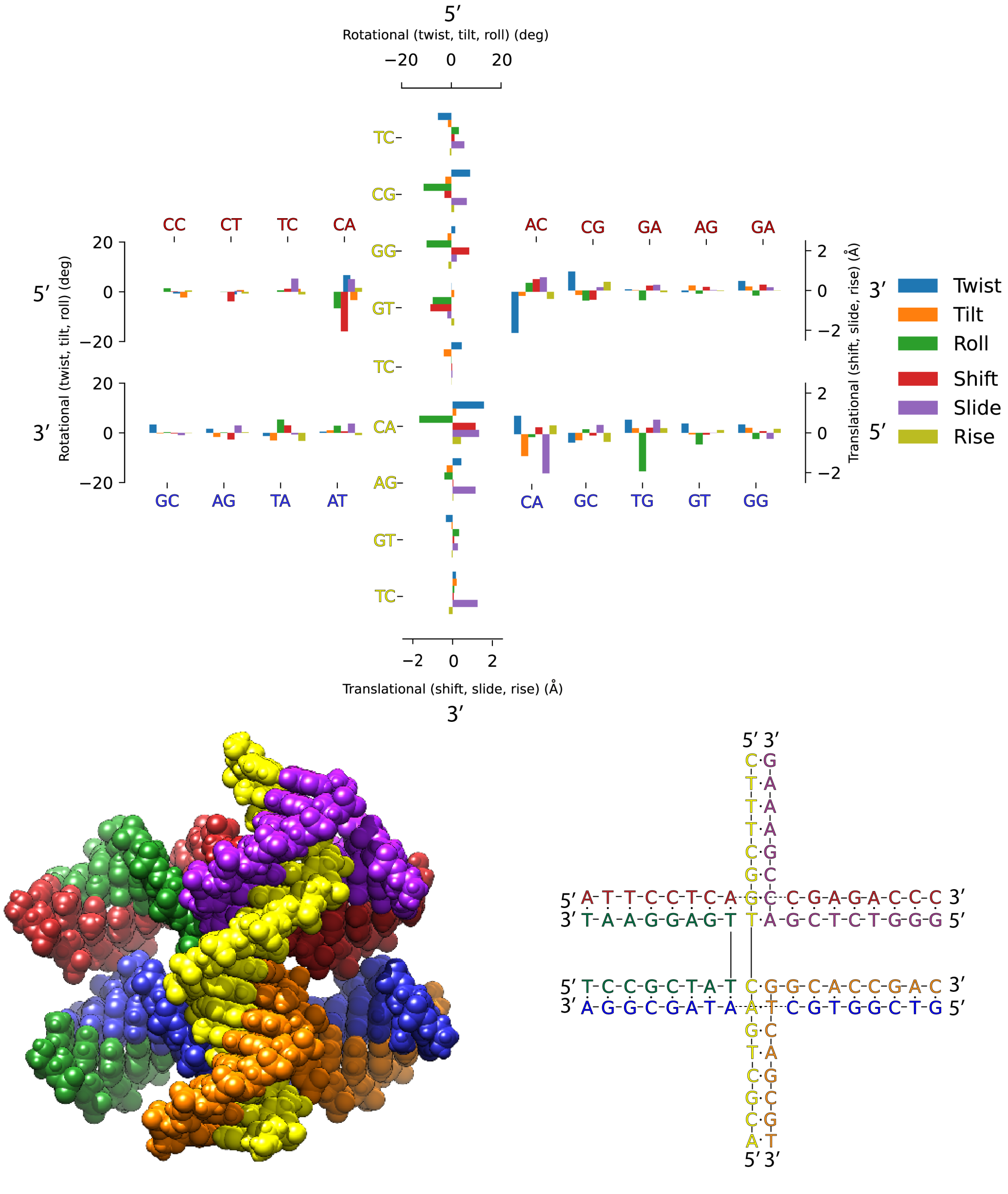

**Figure S17: Average base-pair step deviations of the TO1 isomer from MD**. The graph on top describes the BPS parameter deviations for each duplex from reference B-DNA. The 5' and 3' on each side show the direction and BPS on the continuous strand. Below the graph displays an atomic model of the junction next to a model displaying the sequence.

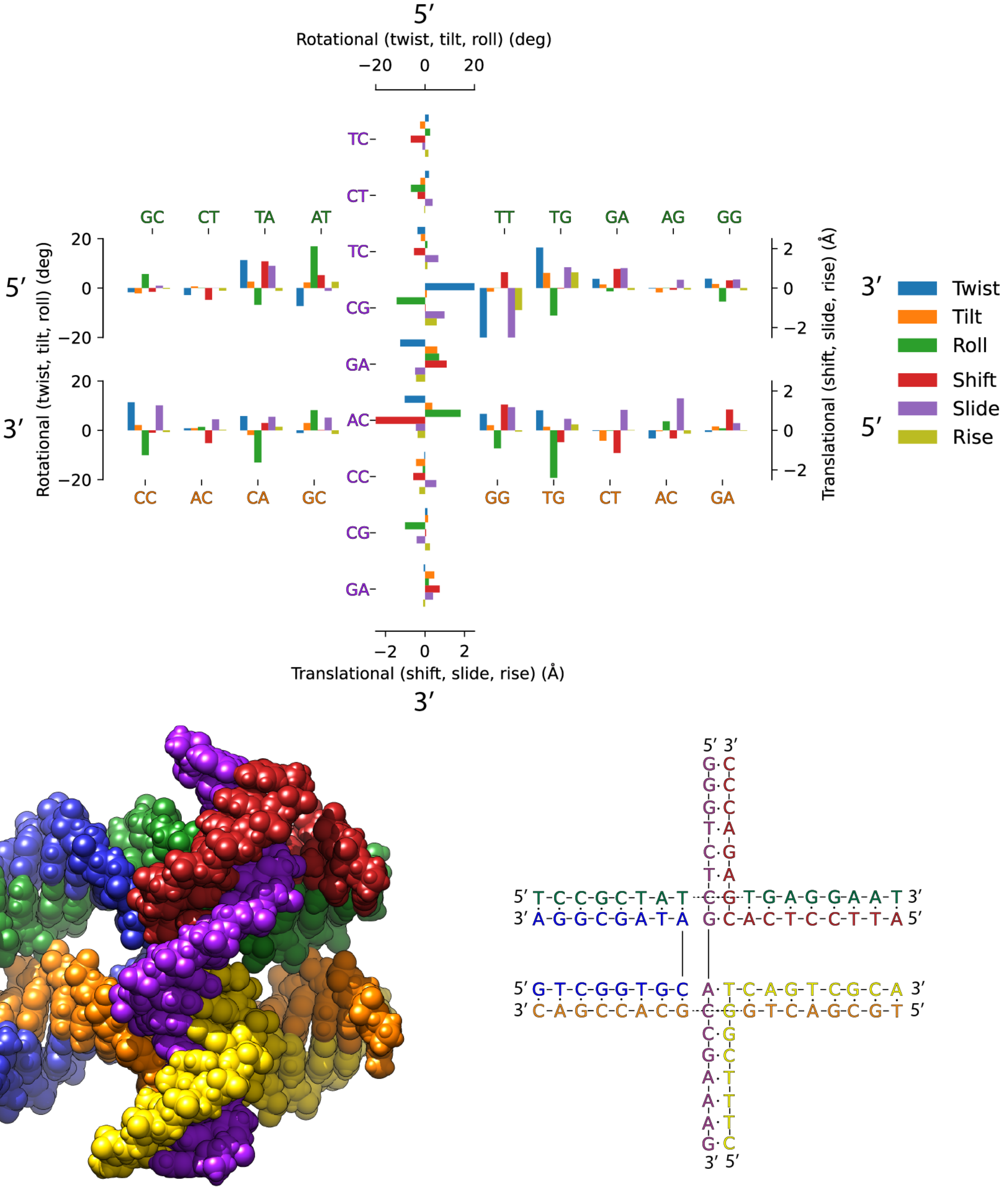


**Figure S18: Average base-pair step deviations of the TO2 isomer from MD**. The graph on top describes the BPS parameter deviations for each duplex from reference B-DNA. The 5’ and 3’ on each side show the direction and BPS on the continuous strand. Below the graph displays an atomic model of the junction next to a model displaying the sequence.

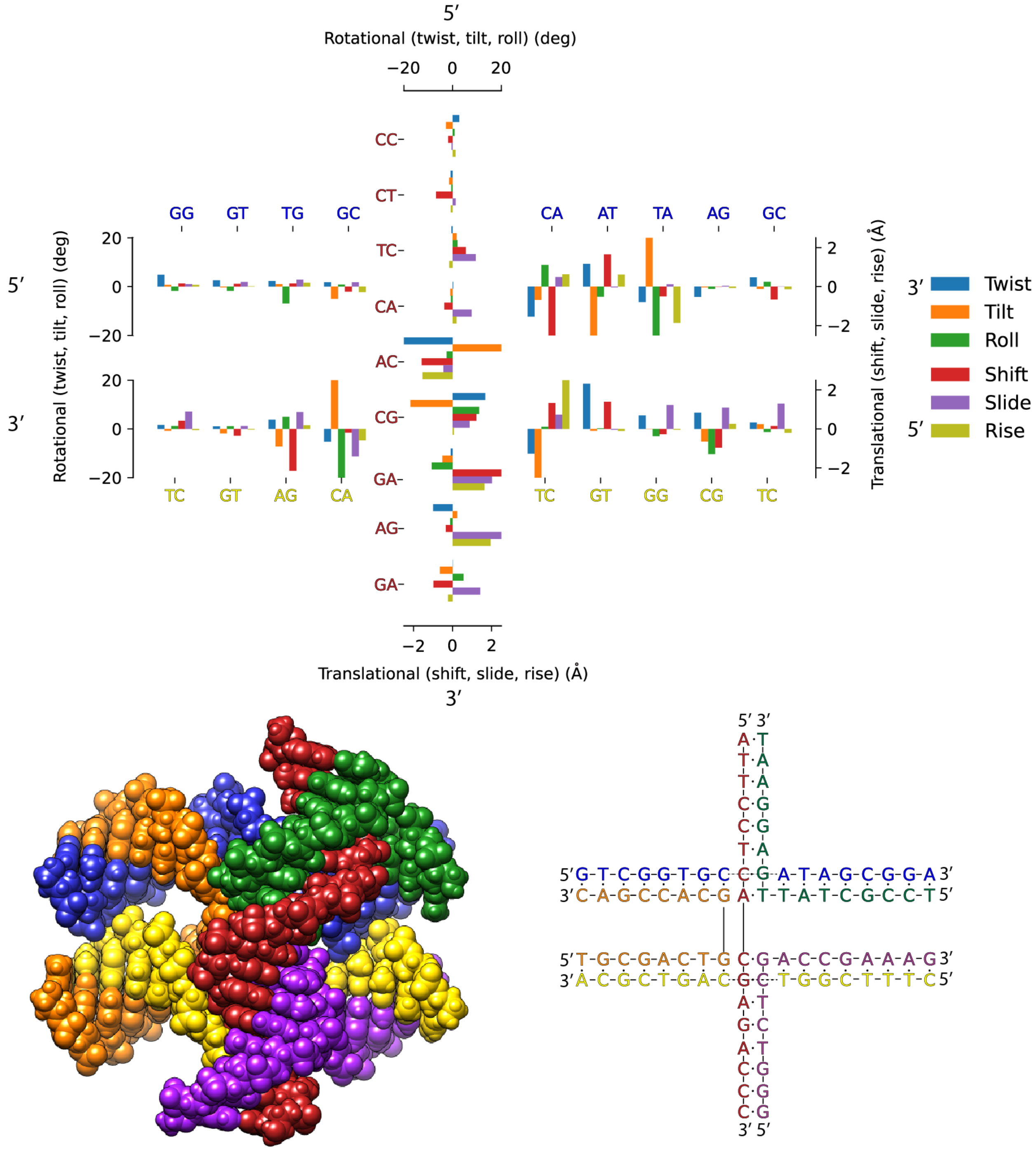


**Figure S19: Average base-pair step deviations of the TO3 isomer from MD**. The graph on top describes the BPS parameter deviations for each duplex from reference B-DNA. The 5' and 3' on each side show the direction and BPS on the continuous strand. Below the graph displays an atomic model of the junction next to a model displaying the sequence.

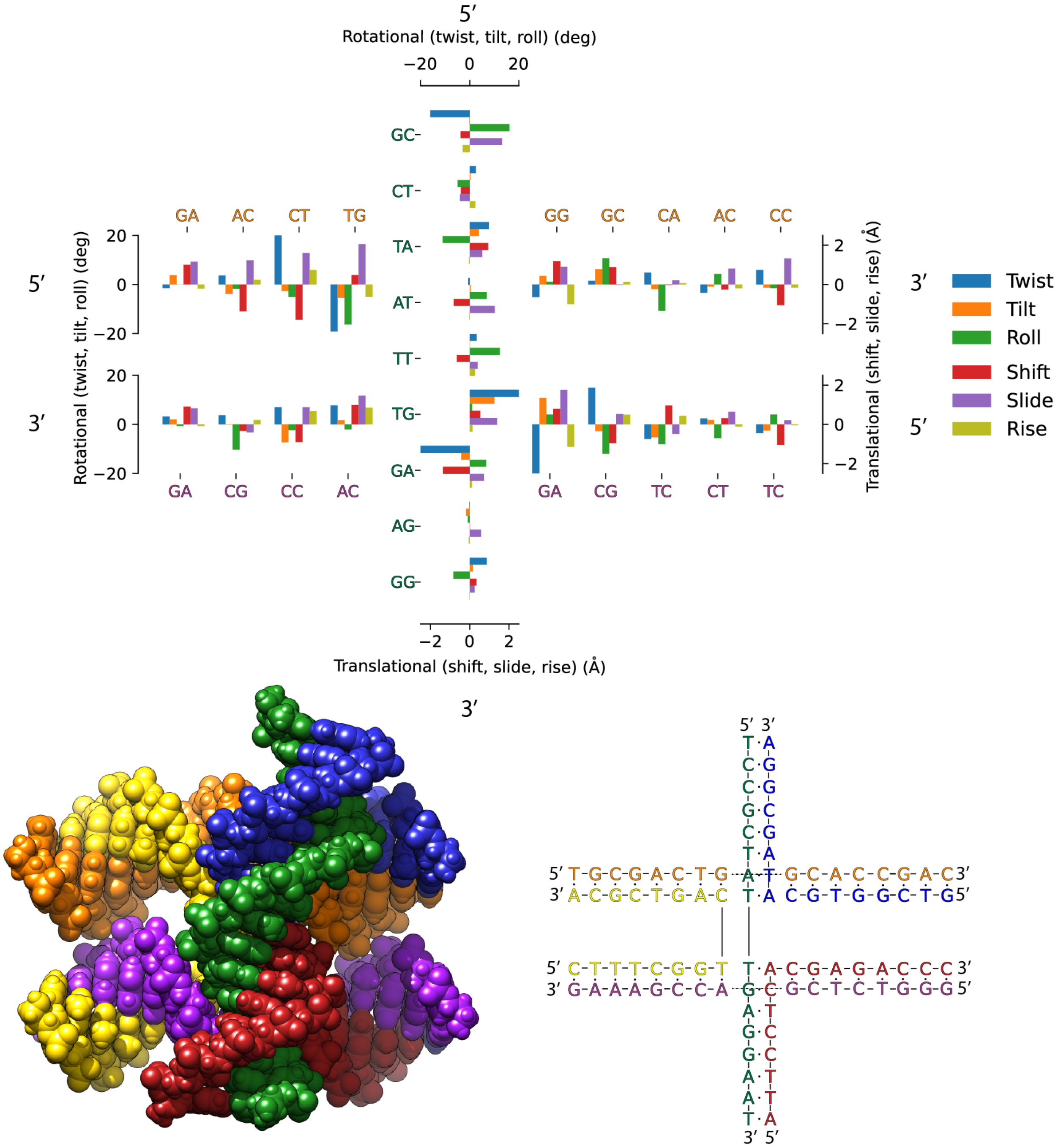


**Figure S20: Average base-pair step deviations of the TO4 isomer from MD**. The graph on top describes the BPS parameter deviations for each duplex from reference B-DNA. The 5’ and 3’ on each side show the direction and BPS on the continuous strand. Below the graph displays an atomic model of the junction next to a model displaying the sequence.

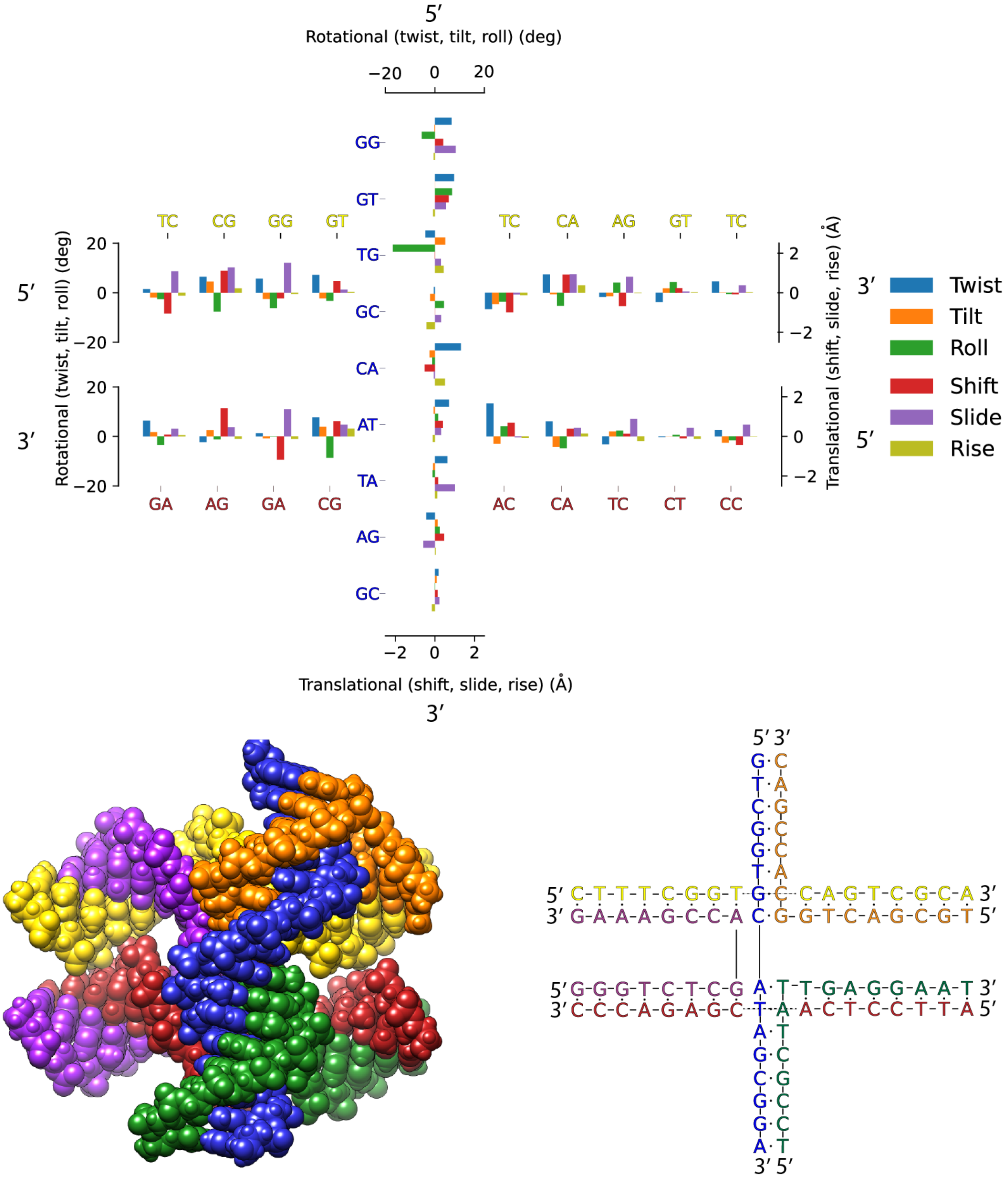


**Figure S21: Average base-pair step deviations of the TO5 isomer from MD**. The graph on top describes the BPS parameter deviations for each duplex from reference B-DNA. The 5’ and 3’ on each side show the direction and BPS on the continuous strand. Below the graph displays an atomic model of the junction next to a model displaying the sequence.

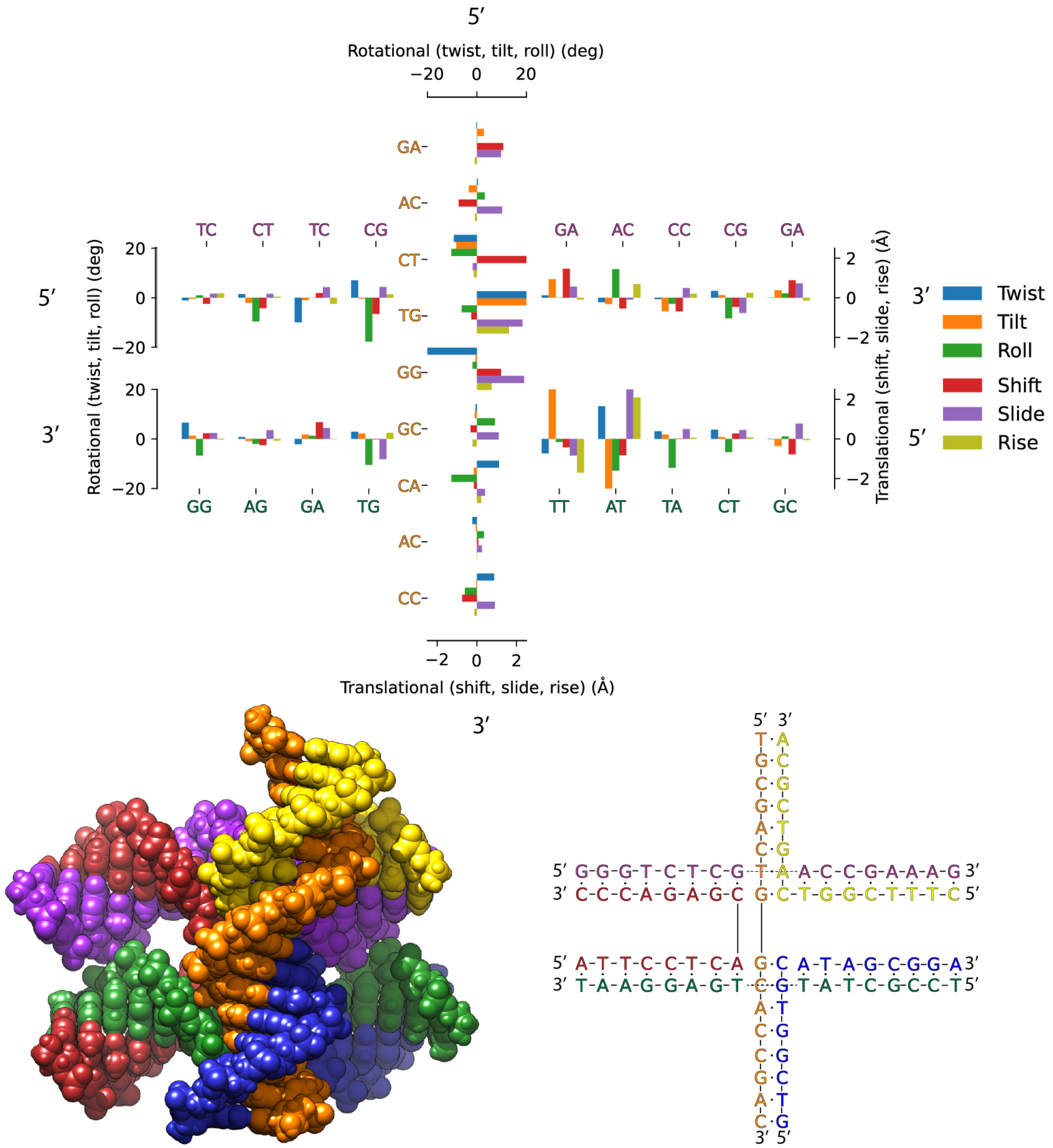


**Figure S22: Average base-pair step deviations of the TO6 isomer from MD**. The graph on top describes the BPS parameter deviations for each duplex from reference B-DNA. The 5' and 3' on each side show the direction and BPS on the continuous strand. Below the graph displays an atomic model of the junction next to a model displaying the sequence.

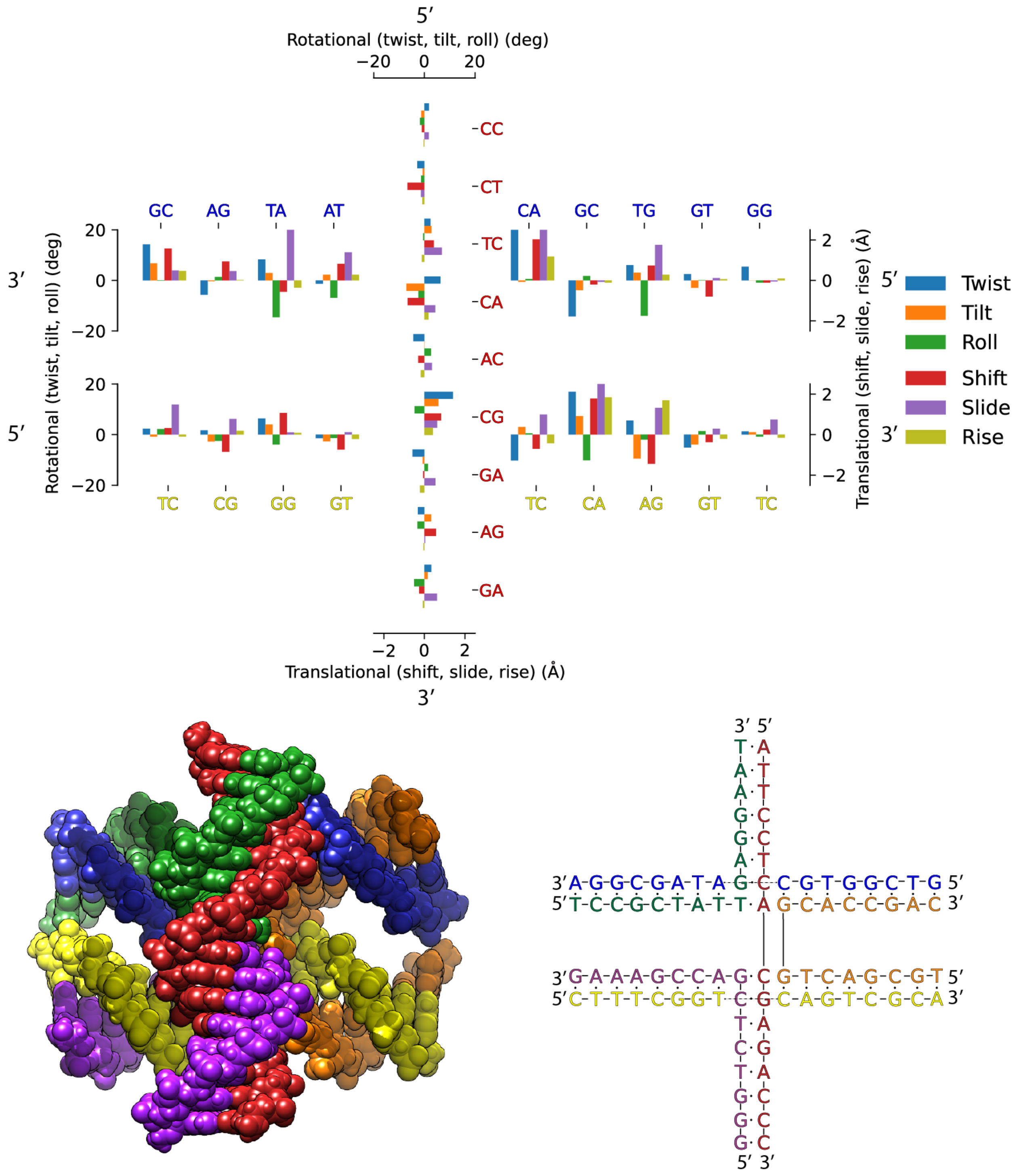


**Figure S23: Average base-pair step deviations of the BO3 isomer from MD**. The graph on top describes the BPS parameter deviations for each duplex from reference B-DNA. The 5' and 3' on each side show the direction and BPS on the continuous strand. Below the graph displays an atomic model of the junction next to a model displaying the sequence.

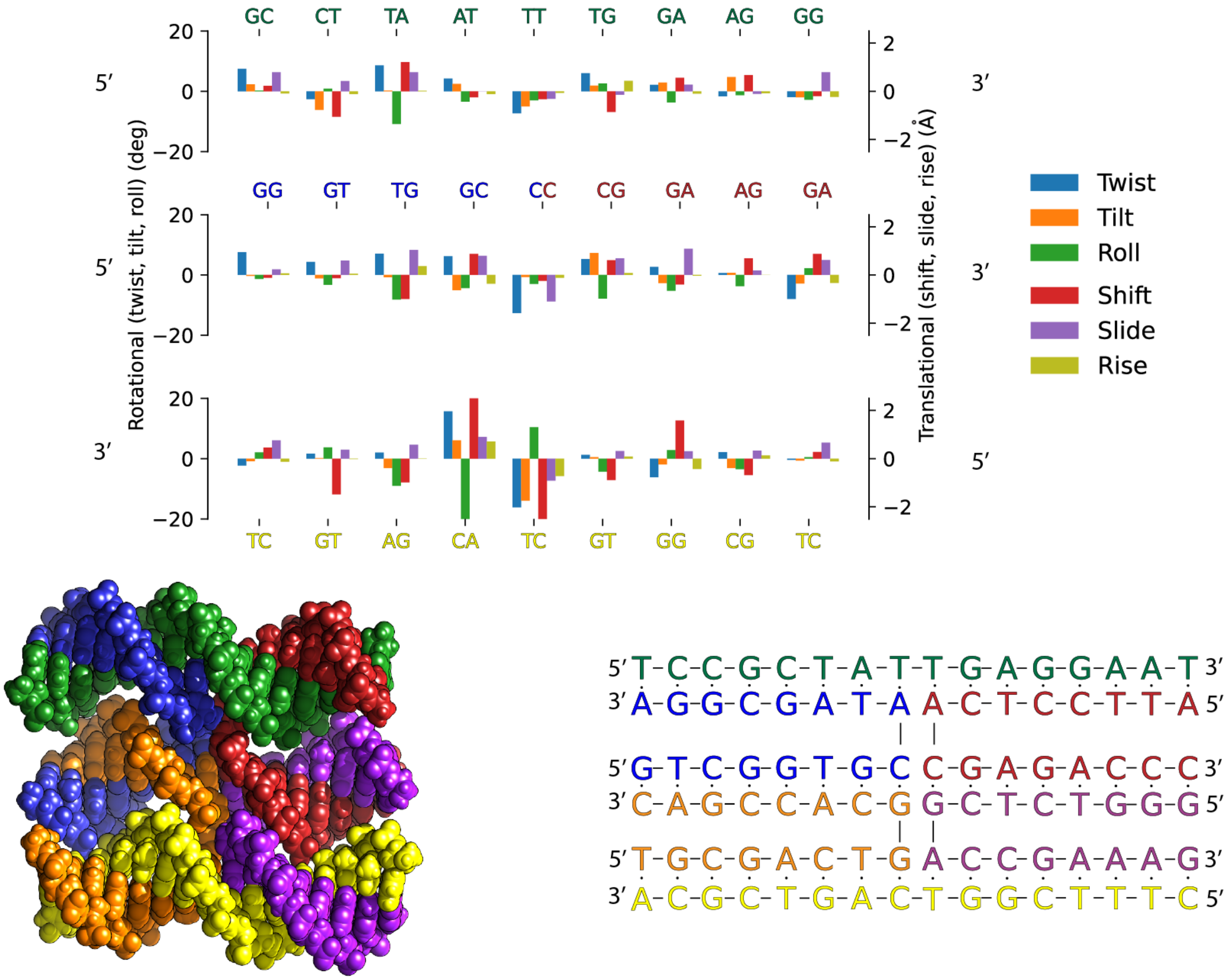


**Figure S24: Average base-pair step deviations of the PP1 isomer from MD**. The graph on top describes the BPS parameter deviations for each duplex from reference B-DNA. The 5' and 3' on each side show the direction and BPS on the continuous strand. Below the graph displays an atomic model of the junction next to a model displaying the sequence.

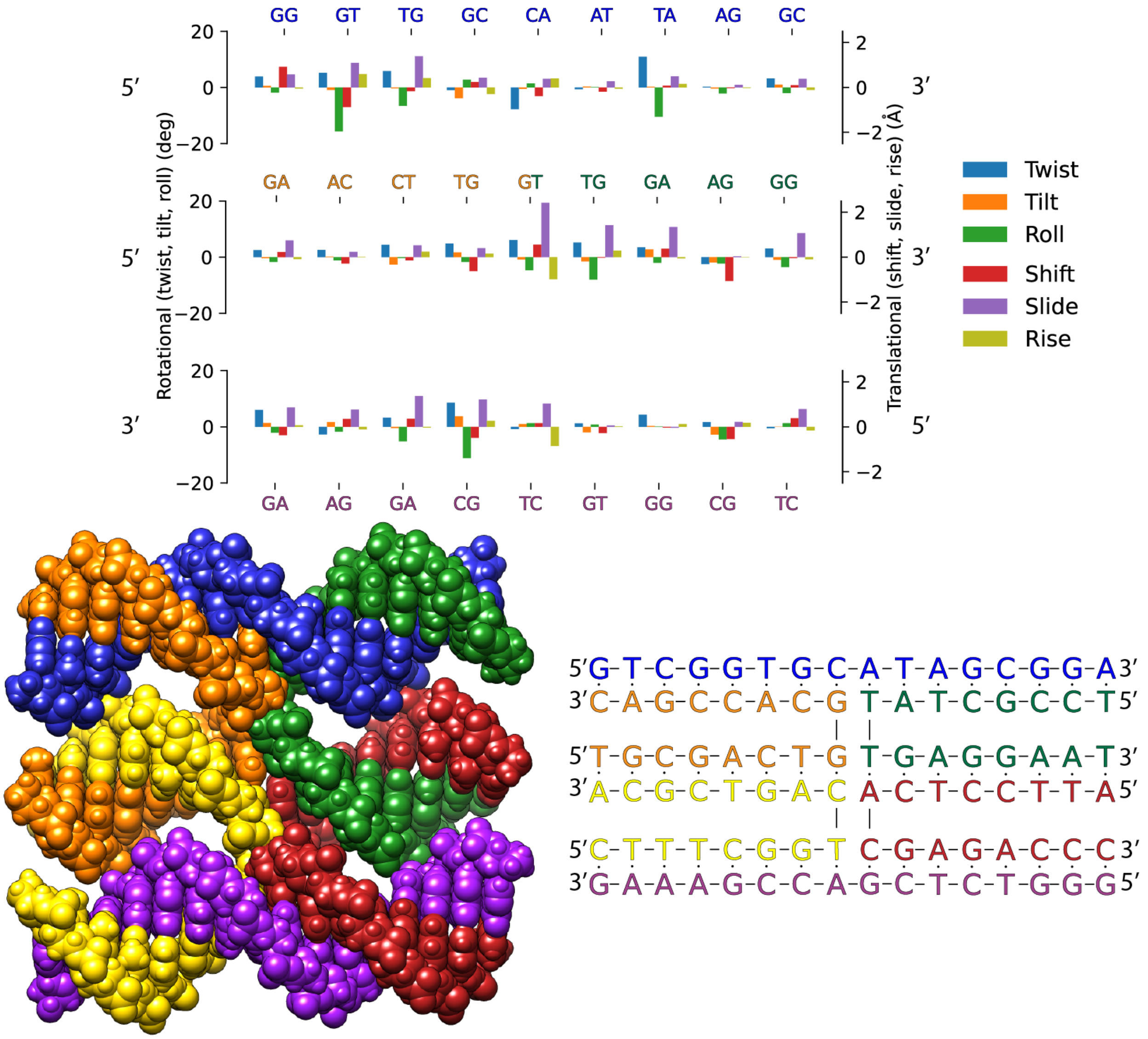


**Figure S25: Average base-pair step deviations of the PP2 isomer from MD**. The graph on top describes the BPS parameter deviations for each duplex from reference B-DNA. The 5’ and 3’ on each side show the direction and BPS on the continuous strand. Below the graph displays an atomic model of the junction next to a model displaying the sequence.

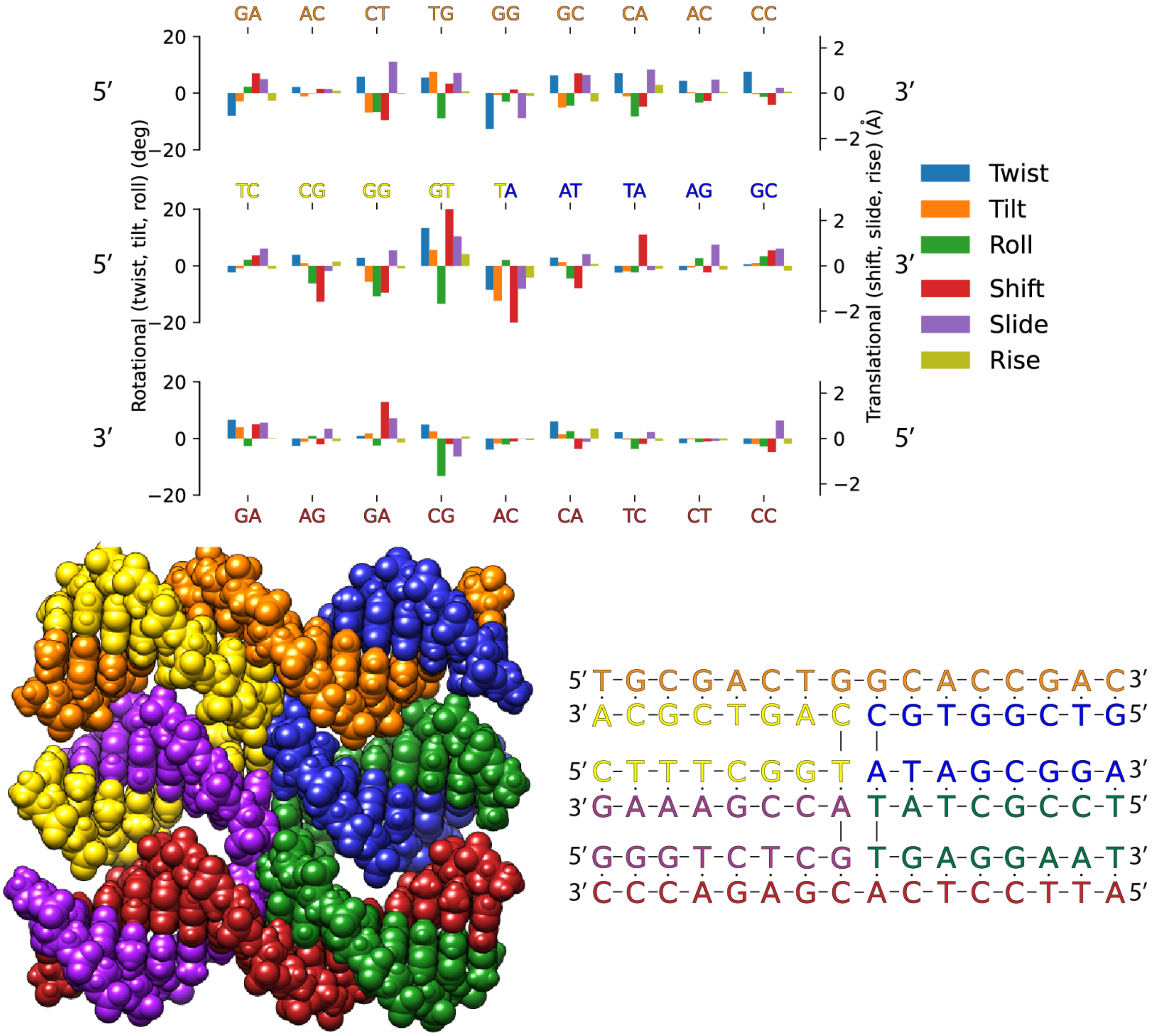


**Figure S26: Average base-pair step deviations of the PP3 isomer from MD**. The graph on top describes the BPS parameter deviations for each duplex from reference B-DNA. The 5' and 3' on each side show the direction and BPS on the continuous strand. Below the graph displays an atomic model of the junction next to a model displaying the sequence.

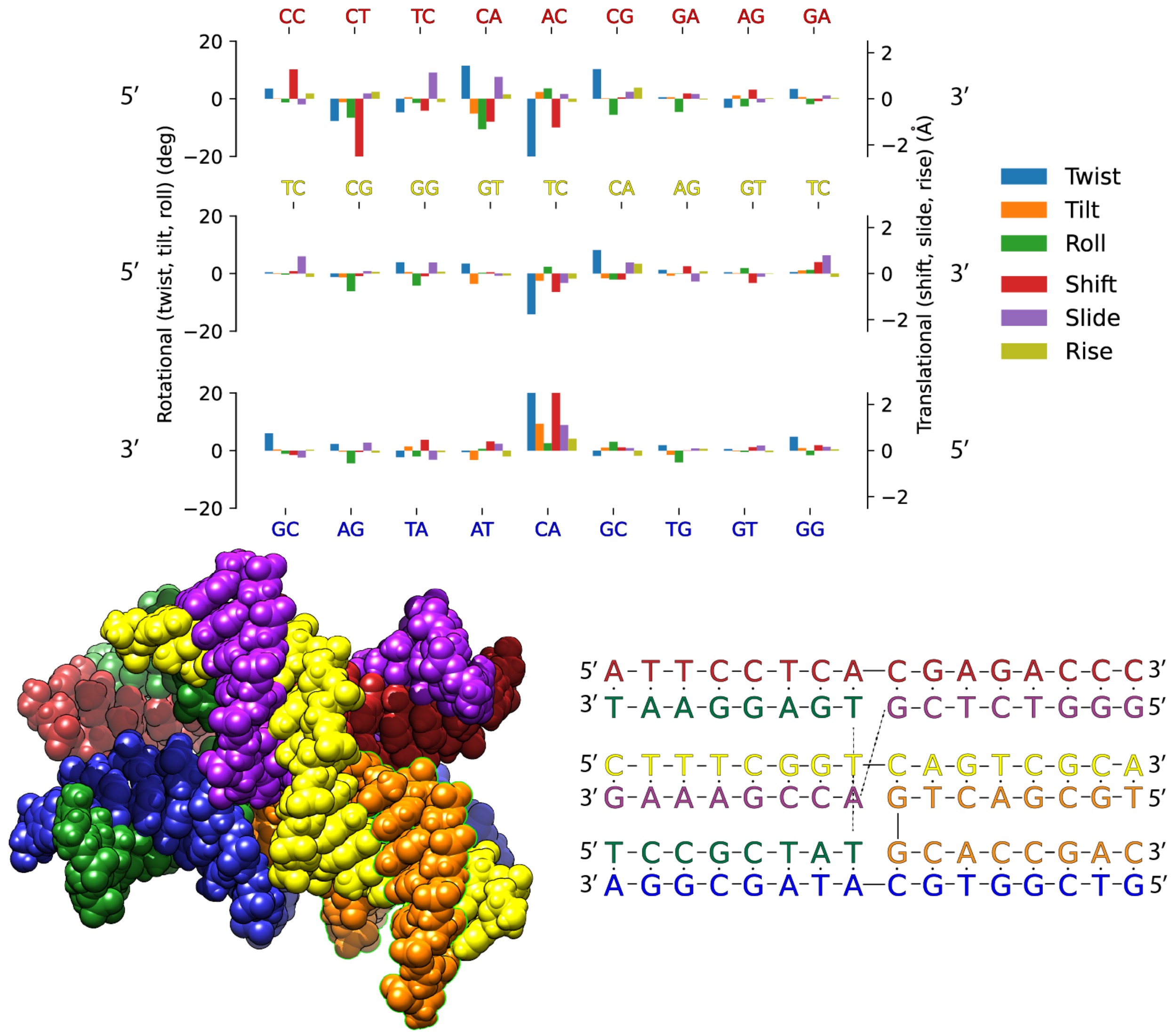


**Figure S27: Average base-pair step deviations of the RT1 isomer from MD**. The graph on top describes the BPS parameter deviations for each duplex from reference B-DNA. The 5' and 3' on each side show the direction and BPS on the continuous strand. Below the graph displays an atomic model of the junction next to a model displaying the sequence.

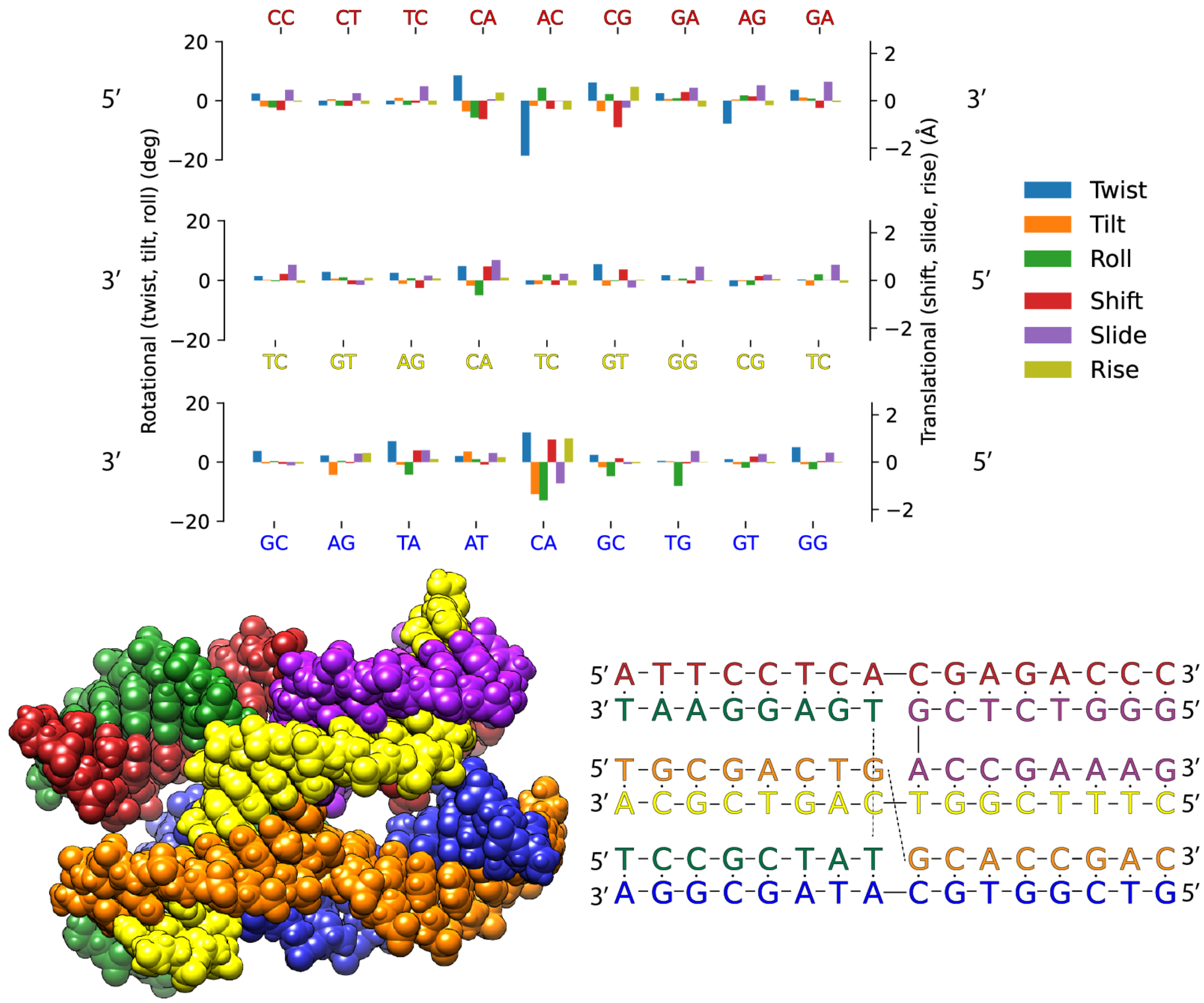


**Figure S28: Average base-pair step deviations of the LT1 isomer from MD**. The graph on top describes the BPS parameter deviations for each duplex from reference B-DNA. The 5' and 3' on each side show the direction and BPS on the continuous strand. Below the graph displays an atomic model of the junction next to a model displaying the sequence.

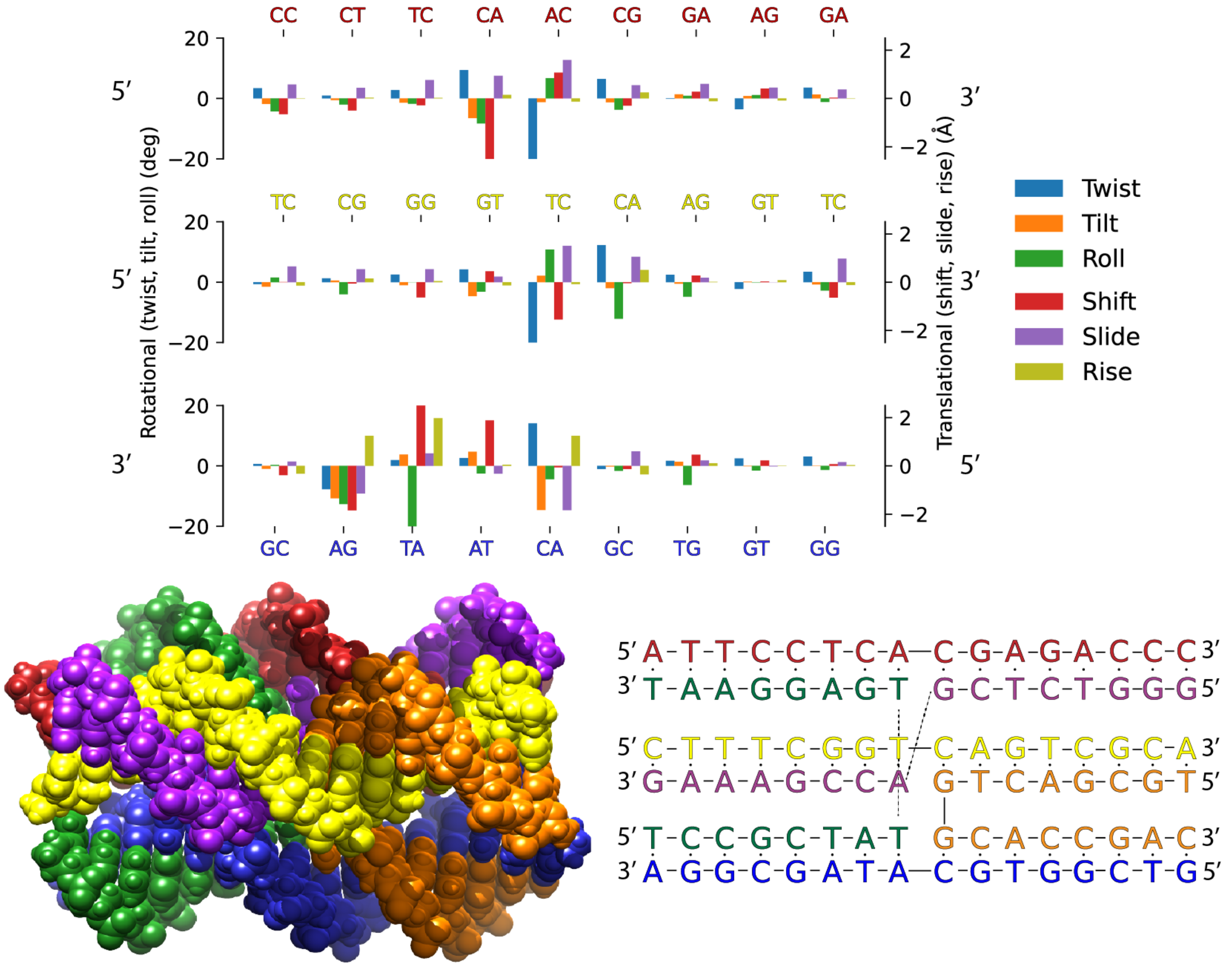


**Figure S29: Average base-pair step deviations of the RP1 isomer from MD**. The graph on top describes the BPS parameter deviations for each duplex from reference B-DNA. The 5' and 3' on each side show the direction and BPS on the continuous strand. Below the graph displays an atomic model of the junction next to a model displaying the sequence.

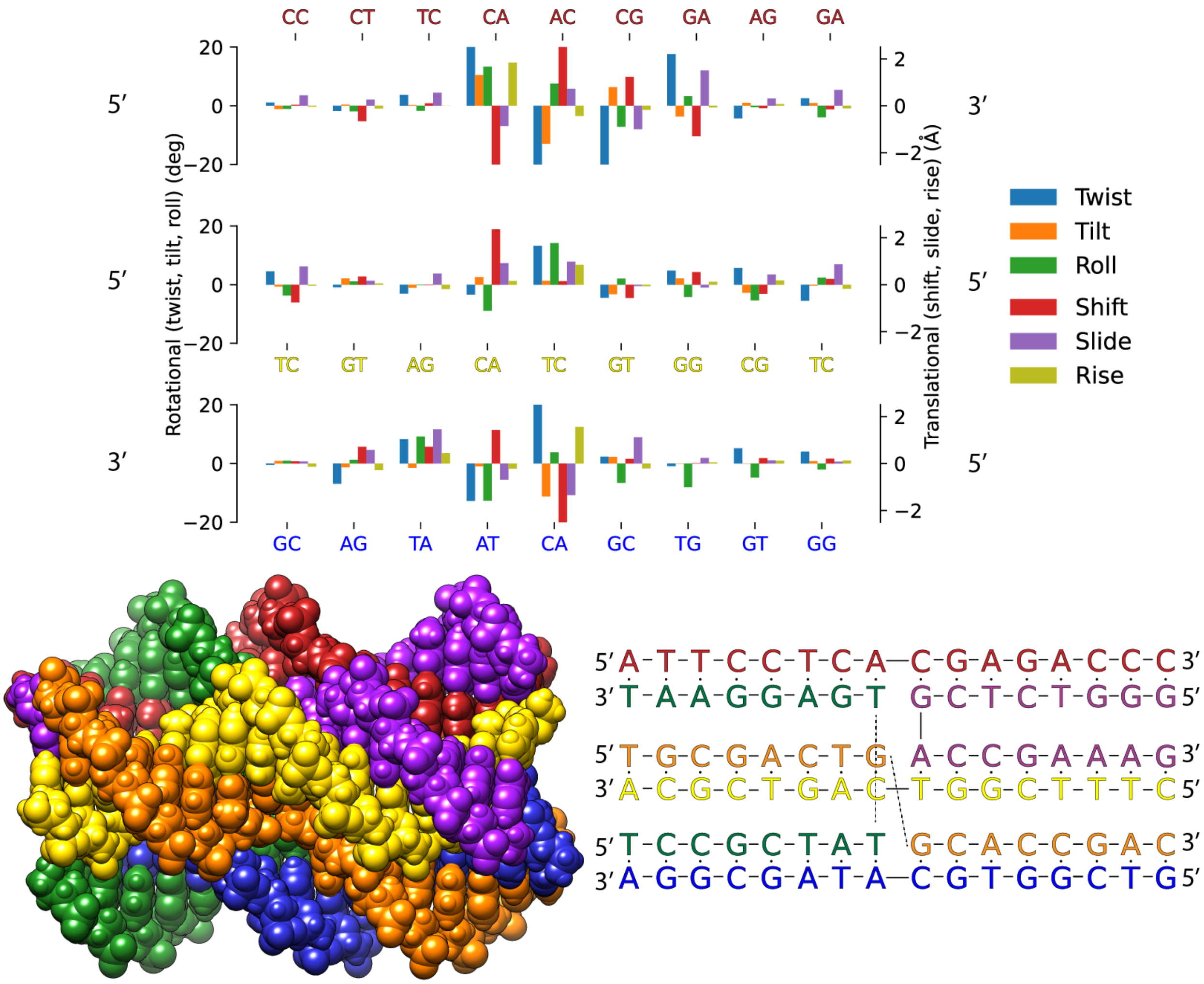


**Figure S30: Average base-pair step deviations of the LP1 isomer from MD**. The graph on top describes the BPS parameter deviations for each duplex from reference B-DNA. The 5' and 3' on each side show the direction and BPS on the continuous strand. Below the graph displays an atomic model of the junction next to a model displaying the sequence.

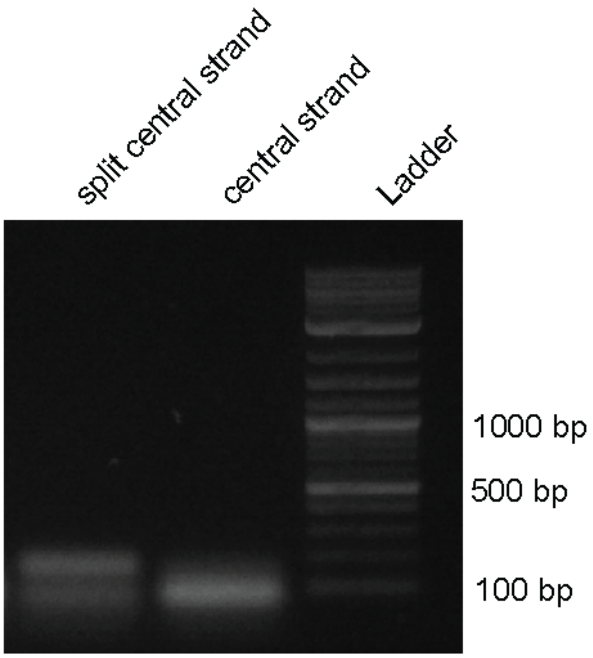


**Figure S31: Electrophoresis gel of central strand routing variants of the tile motif.**

**Table S2: Sequences of the oligonucleotides used to assemble the DNA 6 way junction.**

| **Structure** | **Sequence ID** | **Sequence** | **Length** |
|---|---|---|---|
| DNA 6WJ | 6WJ:1 | *CTTTCGGTCAGTCGCA* | 16 |
| DNA 6WJ | 6WJ:2 | *GGGTCTCGACCGAAAG* | 16 |
| DNA 6WJ | 6WJ:3 | *ATTCCTCACGAGACCC* | 16 |
| DNA 6WJ | 6WJ:4 | *TCCGCTATTGAGGAAT* | 16 |
| DNA 6WJ | 6WJ:5 | *GTCGGTGCATAGCGGA* | 16 |
| DNA 6WJ | 6WJ:6 | *TGCGACTGGCACCGAC* | 16 |

**Table S3: Sequences of the oligonucleotides used to assemble the DNA 6 way junction for the FRET assay.** These sequences were used in combination with the sequences in Table S2 to prepare multiple FRET pair variants. FAM: Fluorescein; TAM: TAMRA.

| Structure | Sequence ID | Sequence | Length |
|---|---|---|---|
| DNA 6WJ-FRET | 6WJ:1-Donor | $FAM - CTTTCGGTCAGTCGCA$ | 16 |
| DNA 6WJ-FRET | 6WJ:2-Acceptor | $TAM - GGGTCTCGACCGAAAG$ | 16 |
| DNA 6WJ-FRET | 6WJ:3-Acceptor | $TAM - ATTCCTCACGAGACCC$ | 16 |
| DNA 6WJ-FRET | 6WJ:5-Donor | $FAM - GTCGGTGCATAGCGGA$ | 16 |
| DNA 6WJ-FRET | 6WJ:6-Acceptor | $TAM - TGCGACTGGCACCGAC$ | 16 |

**Table S4: Sequences of the oligonucleotides used to assemble the tile motif.** The dashes indicate a continuing sequence onto the next line.

| Structure | Sequence ID | Sequence | Length |
|---|---|---|---|
| DNA 6WJ-Tile | 6WJ-Tile:1 | $GTGTGAGGTACCTGCGCAAAACAG -$<br>$- CTGGGGTTTCGA$ | 36 |
| DNA 6WJ-Tile | 6WJ-Tile:2 | $GGCACTAAAGCGCGTCTCGAGTTT -$<br>$- AGGGCCTCACAC$ | 36 |
| DNA 6WJ-Tile | 6WJ-Tile:3 | $ATAGTCCCCAGGCCCTAAACTCGA -$<br>$- GACGCGCTGCAGCCCT$ | 40 |
| DNA 6WJ-Tile | 6WJ-Tile:4 | $AGCGTTCGCCCGAGCGGTCCCTAA -$<br>$- GACGTTAGTGCC$ | 36 |
| DNA 6WJ-Tile | 6WJ-Tile:5 | $TCGAAACCTGCCGTTTGTCTGTAC -$<br>$- CCGTCGAACGCT$ | 36 |
| DNA 6WJ-Tile | 6WJ-Tile:6 | $CACGAGATTGGAACGGGTACAGAC -$<br>$- AAACGGCACCCGCACG$ | 40 |
| DNA 6WJ-Tile | 6WJ-Tile:7 | $GCATACAGCTGGTGCG$ | 16 |
| DNA 6WJ-Tile | 6WJ-Tile:8 | $TACAGGCTCGTCTTAGGGACCGCT -$<br>$- CGGGCTGTATGC$ | 36 |
| DNA 6WJ-Tile | 6WJ-Tile:9 | $CCACAACCAGCCTGTA$ | 16 |
| DNA 6WJ-Tile | 6WJ-Tile:10 | $CGCACCAGTCCAATCTCGTGAGGG -$<br>$- CTGCGGTTGTGG$ | 36 |
| DNA 6WJ-Tile | 6WJ-Tile:11 | $CCTGCGTTTTACTTTT$ | 16 |
| DNA 6WJ-Tile | 6WJ-Tile:12 | $ATTCACGACCTGGGGACTATCGTG -$<br>$- CGGGAACGCAGG$ | 36 |
| DNA 6WJ-Tile | 6WJ-Tile:13 | $GCCGTTCGTCGTGAAT$ | 16 |
| DNA 6WJ-Tile | 6WJ-Tile:14 | $AAAAGTAACCAGCTGTTTTGCGCA -$<br>$- GGTACGAACGGC$ | 36 |
| DNA 6WJ-Tile | 6WJ-Tile:15 | $ATAGTCCCCAGGCCCTAAACTCGA -$<br>$- GACGCGCTGCAGCCCTCACGAGAT -$<br>$- TGGAACGGGTACAGACAAACGGCA -$<br>$- CCCGCACG$ | 80 |
